# Multicellular growth as a dynamic network of cells

**DOI:** 10.1101/2023.11.02.565242

**Authors:** Piyush Nanda, Julien Barrere, Thomas LaBar, Andrew W. Murray

**Author notes:** Lead contact (A.W.M.).

## Abstract

Cell division without cell separation produces multicellular clusters in budding yeast. Two fundamental characteristics of these clusters are their size (the number of cells per cluster) and cellular composition: the fractions of cells with different phenotypes. However, we do not understand how different cellular features quantitatively influence these two phenotypes. Using cells as nodes and links between mother and daughter cells as edges, we model cluster growth and breakage by varying three parameters: the cell division rate, the rate at which intercellular connections break, and the kissing number (the maximum number of connections to one cell). We find that the kissing number sets the maximum possible cluster size. Below this limit, the ratio of the cell division rate to the connection breaking rate determines the cluster size. If links have a constant probability of breaking per unit time, the probability that a link survives decreases exponentially with its age. Modeling this behavior recapitulates experimental data. We then use this framework to examine synthetic, differentiating clusters with two cell types, faster-growing germ cells and their somatic derivatives. The fraction of clusters that contain both cell types increases as either of two parameters increase: the kissing number and difference between the growth rate of germ and somatic cells. In a population of clusters, the variation in cellular composition is inversely correlated (r^2^=0.87) with the average fraction of somatic cells in clusters. Our results show how a small number of cellular features can control the phenotypes of multicellular clusters that were potentially the ancestors of more complex forms of multicellular development, organization, and reproduction.

## Introduction

The origin of multicellularity was a major transition in Earth’s life history^1^. One difficulty in this transition was the need for new reproductive strategies that allow faithful inheritance of multicellular characteristics as opposed to simple cell division in unicellular organisms^2–4^. Unicellular organisms can evolve multicellularity either by colliding and sticking together (aggregative multicellularity) or by failing to separate from each other after cell division (clonal multicellularity)^5–7^. Two simple characteristics of the initial multicellular forms were their size (the number of cells in a cluster) and composition (the proportion of different cell-types) but their extinction or evolution makes it difficult to understand how these features evolved. Even in series of phylogenetically related organisms that differ in their degree of multicellularity, like the *Volvocales*, the original unicellular and multicellular forms no longer exist.

One alternative to comparative approaches is to create and investigate multicellularity in the lab by using genetic engineering^8^ or experimental evolution. Recently, the budding yeast, *Saccharomyces cerevisiae*, has emerged as a model organism for understanding aspects of multicellularity^8–10^, including cluster size and cluster composition (in clusters with multiple cell types). Selecting for faster settling^9^ or production and consumption of public goods^10^ leads to the evolution of clonal multicellularity. Cluster size is under selection when faster sedimentation^11^ or public goods sharing^8^ provide a fitness advantage, while cluster composition affects fitness when division of labor or metabolic interactions provide an advantage^12^. Understanding the parameters that control cluster size and composition will offer insights into the evolution of simple multicellularity.

The growth and division of multicellular clusters results from the balance between cell division, which adds cells, and the fragmentation of clusters due to chemical processes and physical forces^13^ that break the linkage between cells in the cluster. For clusters that contain multiple cell types, the transition rates between different cell types and their proliferation rates control cluster composition^614^. Previous work has highlighted genotypes that control maximum cluster sizes and internal physical forces. For example, changes in cell shape^9^ or the strength of cell-cell connections^15^ affect cluster size. In budding yeast, the failure to promptly dissolve the primary septum, the portion of the cell wall that links daughter cells to their mothers, leads to multicellularity. Experimental evolution of multicellularity has resulted in mutations in genes that affect the linkage between the mothers and daughters (*ACE2, CTS1, GIN4*) or control cell shape (*CLB2, ARP5*)^10^ but we do not know how these mutations affect cluster growth and fragmentation to produce a steady-state distribution of cluster sizes. Modelling has explored the effect of geometric constraints on individual clusters ^11^ and suggests that changing cell shape or the geometry of the linkages to a single cell increases cluster size more than increasing strength of the linkages between individual cells^16^.

While these studies discuss the parameters that set fundamental limits on cluster size, the underlying phenomena that control the mean and distribution of cluster sizes remain unclear. An ideal model would enable *ab initio* prediction of cluster size and its distribution from a few measurable variables that can be directly linked to gene products or pathways in cells. Such predictions can be useful in engineering clusters of desired sizes or understanding the behavior of multicellular clusters in different environments where underlying variables (like rates at which cells grow or dissolve their linkages to other cells) differ. For example, how does a change in budding pattern in yeast alter the cluster size distribution? How is this distribution affected if the growth rate of cells changes? In clusters where differentiation has produced a division of labor between different cell types^12,17,18^, the distribution of cluster compositions will depend not only on the transition rates between different cell types and their rates of proliferation but also on the cluster size distribution. How does the cluster size distribution change the composition of such primitive, differentiated clusters?

We studied the size and composition of multicellular cluster by constructing a dynamic network model that represents cells as nodes and the connections between mothers and daughters as edges (**Figure 1A**). Using simulations and experiments, we analyzed the effect of varying three parameters: the cell division rate, the rate at which links between cells break, and the kissing number (the maximum permissible number of connections to one cell). We show that a model with a steric limit on the kissing number and links that are more likely to break as they age best captures the experimentally observed distribution. The faster cells divide relative to the link breaking rates, the more closely cluster sizes approach the upper limit set by the kissing number. In differentiating clusters with germ and somatic cells, increasing the kissing number and optimizing the difference in the division rate between the two cell types maximizes the fraction of clusters that contain both cell types. We discuss the implications of our work for understanding the evolution of adaptive multicellular phenotypes and extensions of this model to capture complex multicellular properties.

**Figure 1.**
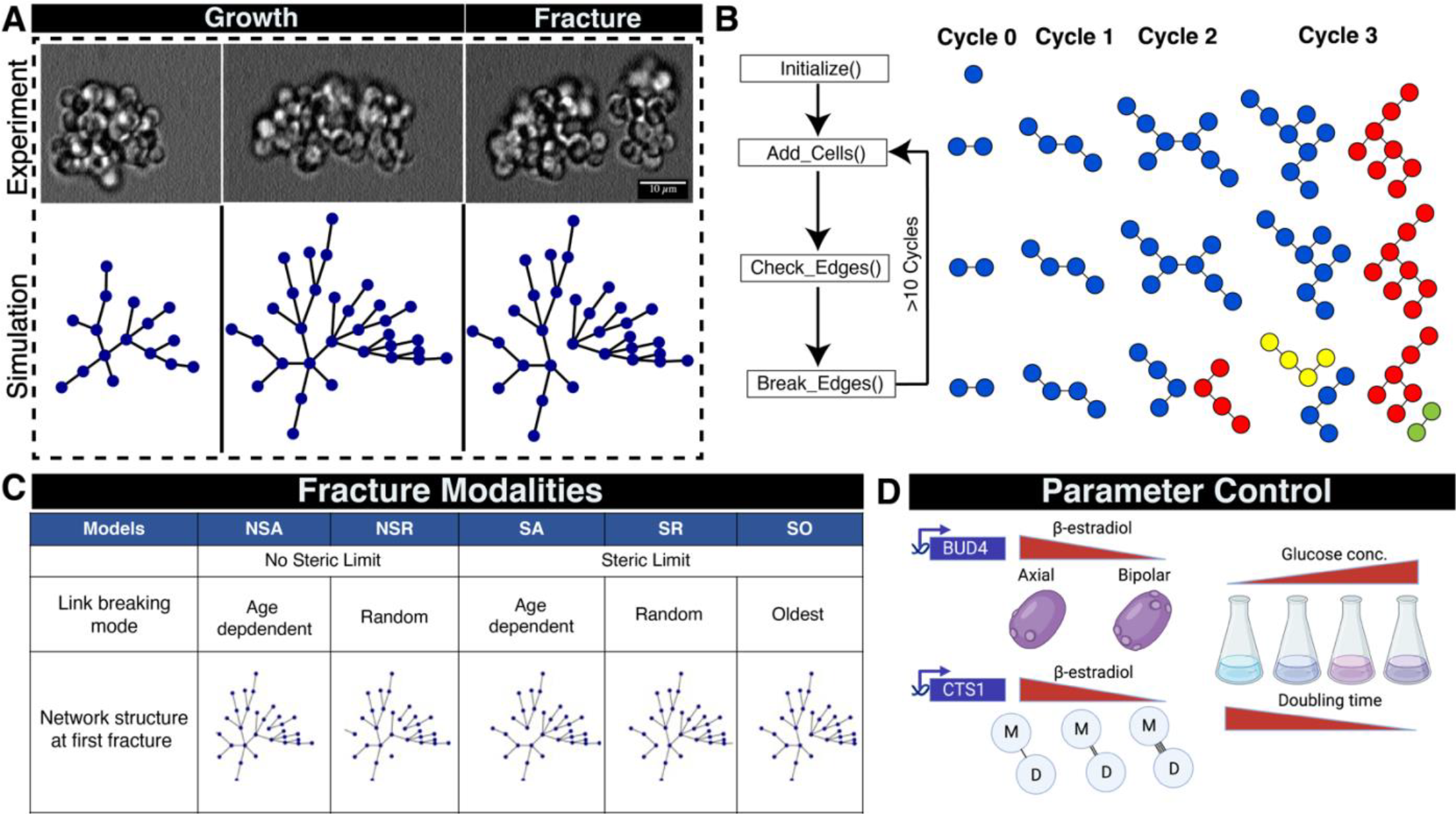
A dynamic network-based approach to examine the mode of breaking in yeast multicellular clusters. A) Three sequential images showing the growth and fracture of a multicellular yeast cluster over several generations and a simulation using the SA dynamic network model in which each blue dot is a cell and each line is a link between mother and daughter cells. B) Illustration of the dynamic network model to simulate multicellular growth and fracture. Different node colors indicate physically separate clusters. C) Five models of fracture based on whether they use a steric limit and how the links between cells break. D) Experimental approaches to regulate three parameters: the mode of budding (kissing number), link breaking rate, and doubling time

## RESULTS

### A dynamic-network approach to model the growth of multicellular clusters

We programmed a dynamic network model to simulate the growth and breakage of clusters and predict steady-state distributions of cluster sizes (**Figure 1B**). In a dynamic network, the edges and nodes can change (appear or disappear) over time. We represent cells as nodes and their connections as edges. Each network represents an individual cluster and grows as cells divide, with division rate (*λ*), and fragments at a specific edge when a certain set of conditions are met. Each time a cell divides, it creates a new link to a daughter. We subject the dynamic network to several cycles of growth and fracture to reach steady state distribution (**Figure 1B**). Increasing crowding around a mother cell leads to the development of steric forces due to geometric constraints, which we capture as the kissing number, *κ*, the maximum number of links a cell may have to its daughters. The concept of kissing number is derived from geometric limits on the packing of spheres^19^ i.e., the maximum number of spheres that can surround a given sphere. Once the number of links to a cell approaches the kissing number, one or more cell-cell connections start to fracture. The fracture can either happen probabilistically, i.e., each connection has a probability of breaking based on an underlying probability distribution, or deterministically, for example, the oldest connection breaks. The fracture partitions the cells in the parental cluster into two new clusters, with the ratio of their sizes determined by which link breaks. We allow cluster growth and fragmentation to continue until the distribution of cluster sizes in a population reaches a steady state. In our model, the cell division rate, the rules for breaking links, and the kissing number can be varied independently, allowing us to explore multiple modalities with different rules for cluster fragmentation.

We began asking if all cluster sizes in a yeast population can grow and proliferate to produce the full distribution of cluster sizes. We tested whether clusters of different sizes, selected from the same distribution can give rise to the parental distribution. Using a *cts1Δ* strain, which makes clusters of 3-10 cells because it lacks the chitinase (Cts1p) that helps to degrade the primary septum^12^, we used Fluorescent Activated Cell Sorting (FACS) to separate the population into four quartiles (Q1, Q2, Q3 and Q4), by cluster size, allowed them to proliferate independently for 48 hours, and measured the distribution of cluster sizes (**Figure S1A**). All four populations recapitulate the original distribution, strongly suggesting that it is generated from an underlying process whose parameters are similar across the population.

We considered five possible modalities of cluster fracture (**Figure 1C**) and modeled their effect on the cluster size distribution. There are two types of constraint that could limit cluster size: i) Steric limited (S), the clusters grow and reach a packing limit at which steric forces fracture the cluster to relieve the strain. ii) Non-Steric limited (NS), the links between cells decay over time, and the clusters fracture because links are chemically degraded rather than because any of its cells exceed the kissing number. The models are then subdivided by specifying the rules that determine which link breaks to fracture a cluster. One choice is deterministic: in the SO (steric, oldest), the oldest link in the entire cluster breaks, separating the two cells in the cluster with the highest number of links as the older of these cells has reached the maximum number of cells it can be surrounded with. Alternatively, a stochastic process assigns a fracture probability to each link. In one pair of models, once the maximum number of connections per cell (SR) or breaking time (NSR) is reached, one or more randomly chosen link fractures. Finally, if the chronological age (A) of the link matters, the longer it has been present in the cluster, the more likely it is to have fractured. In these models (SA or NSA), the probability that a link breaks per unit time is a constant given by the link breaking rate, *δ*, and the probability that a link survives declines exponentially with its age: *p*(survival to time *t*) = exp(-*δt*). In our simulations, each link is assigned a randomly chosen survival time from this exponential distribution. Note that when a cluster breaks into two new clusters, many of the links are already several generations old, and are thus more likely to break in the future.

We set out to test which of the models best explains experimental cluster size distributions. We engineered strains where we can control the biological equivalents of the three model parameters: the cell division rate (*λ*), the kissing number (*κ*) and the link-breaking rate (*δ*). We altered the kissing number by controlling the expression of *BUD4*^20^ (**Figure 1D**) in a strain deleted for *ACE2* to switch the budding pattern from axial (*κ* = 5, *BUD4* ON) to bi-polar (*κ* = 8, *BUD4* OFF). We controlled the link-breaking rate (*δ*) by tuning the expression of the chitinase gene (*CTS1*) and controlled the cell division rate (*λ*) by altering the concentration of the carbon source in the media^21^. Finally, we compared the simulated cluster size distribution obtained from all the models with the experimental distribution using an imaging pipeline that determines the cluster size distribution (**Figure S2, S3 and S4**).

### Steric-limited age-dependent fracture (SA) based dynamic network model predicts cluster size distributions

To test model predictions, we first varied the kissing number parameter (*κ*) while keeping the other two parameters (*δ* and *λ*) constant. We changed the levels of Bud4p protein in an *ace2*Δ strain by using a β-estradiol-inducible promoter^22^. Bipolar budding (*BUD4*-OFF) allows a greater number of daughter cells to remain attached to mother cells before the onset of steric forces as buds emerge from opposite poles. We confirmed the axial and bipolar budding patterns by imaging calcofluor white-stained clusters (**Figures 2A and 2D**). Using our imaging pipeline (**Figure S2, S3 and S4**), we obtained the cluster size distributions corresponding to *BUD4*-ON and *BUD4*-OFF. For *BUD4*-OFF, the maximum number of cells in a cluster ranged between 128 and 256. This implies a kissing number of 8, as exceeding this would produce clusters with more than 256 cells. Meanwhile, the maximum number of cells per cluster for *BUD4*-ON clusters was between 16 and 32, implying a kissing number of 5. The cell division rate (*λ*) of the clusters was estimated from growth curves, and the link breaking rate (*δ*) for the SA model was set to a very low value (*δ* =0.01*h*^−1^) such that link breaking does not limit the size of the clusters. In simulations, the number of growth cycles was sufficient to allow the simulated cluster size distributions reach a steady state (**Figure S6E and S6F).**

**Figure 2.**
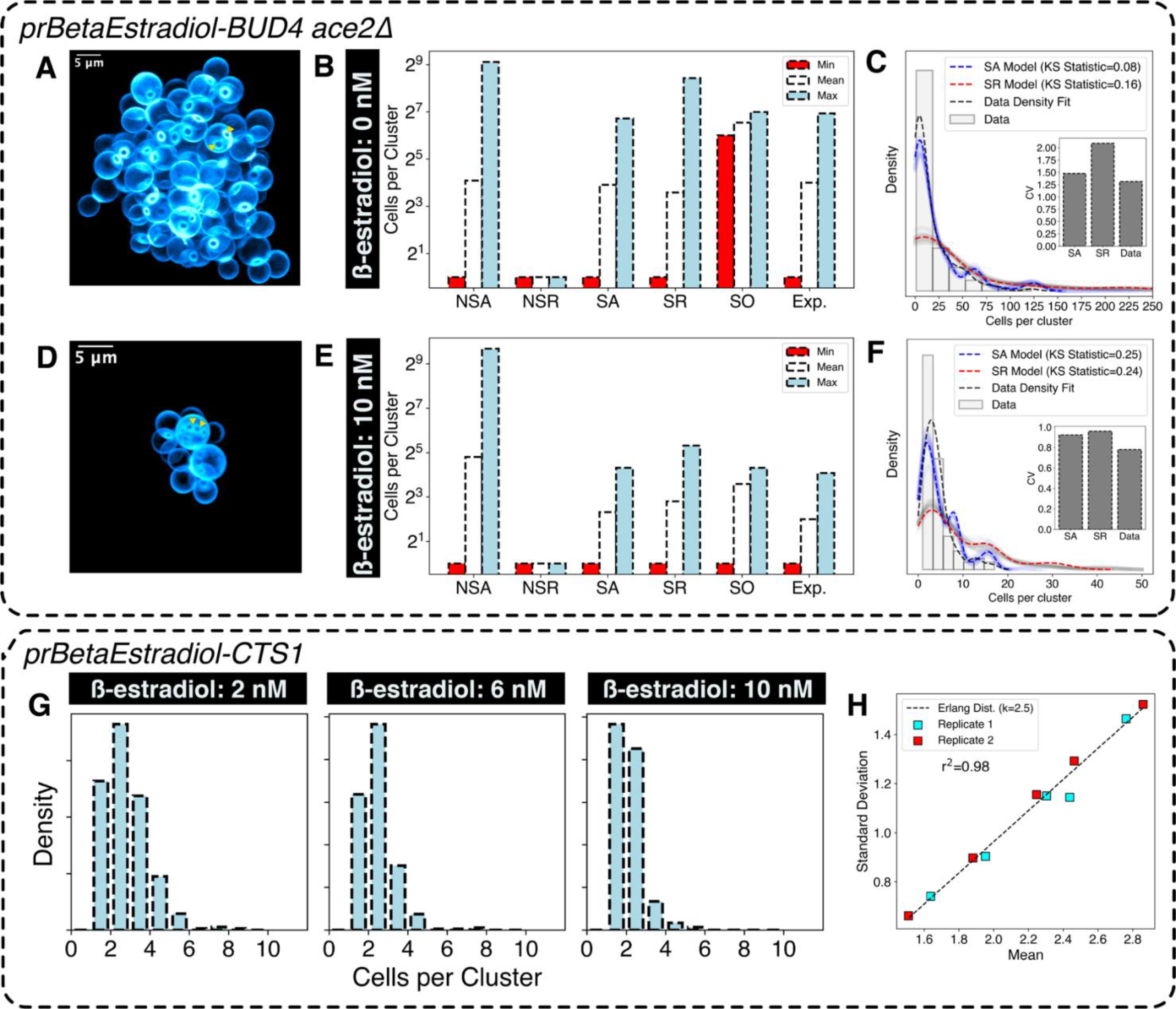
Sterically limited, age-dependent (SA) model recapitulates experimental cluster size statistics. A) -dimensional CLSM (Confocal Laser Scanning Microscope) rendering of a multicellular cluster with uninduced UD4 expression showing the location of bud scars (The triangles indicate bud scar location) B) Cluster size istribution statistics for different models and experimental data (Exp.) for high kissing number (*κ* =8). C) istribution obtained from models (SA and SR) and experimental measurements for uninduced BUD4 expression n=109 clusters). The faint lines show simulation repeats with independent seed values, the darker lines show their average. D) 3-dimensional rendering of multicellular clusters with fully induced BUD4 expression (The triangles ndicate bud location) E) Cluster size distribution statistics for different models and experimental data for low kissing umber (*κ* =5) F) Distribution obtained from models (SA and SR) and experimental measurements for full induction n=114 clusters). The faint lines show simulation repeats with independent seed values, the darker lines show their average. G) Cluster size distribution for different levels of Cts1p controlled through β-estradiol induction. H) Relation between mean and standard deviation of cluster size distribution obtained in (G) for two independent biological replicates (n≈400 clusters per condition). The linear regression fit is to an Erlang’s distribution with shape parameter=2.5.

When comparing experimental data and simulations for the two different kissing numbers (*BUD4*-OFF and *BUD4*-ON) the SA model best predicted the cluster size distributions in both cases (**Figure 2B, C, E and F**). The correlated increase between cluster size and kissing number strongly suggests that cluster size is steric-limited.

None of the other models could accurately fit the experimental data (**Figure S6A, S6B and S6C**). The SR model, which assigns an equal probability of breaking to all the edges, produces a cluster-size distribution that has a higher maximum value than experimentally seen: breaking the oldest edge produces daughter clusters of equal sizes, whereas breaking the newest link produces highly asymmetric daughter clusters leading to a coefficient of variation (CV) in cluster size larger than the experimental data (**Figure 2C and 2F**). All the NS models, which lack a steric component, fail to explain the difference in cluster size between the *BUD4*-ON and *BUD4*-OFF data (**Figure S6A and S6C**). For example, Model NSA produces some clusters that are larger than those seen in any experimental condition (>500 cells per cluster). Model NSR produces a cluster size distribution restricted to 2-6 cells per cluster. Model SO produces a bimodal distribution of cluster sizes corresponding to 128 and 64 cells per cluster. Breaking at the oldest node will always produce clusters of equal sizes as the cluster is symmetric about the oldest edge. Therefore, the clusters would have 128 cells per cluster when they are about to break or 64 cells per cluster after they break.

One potential drawback of our modeling strategy is that all the cells in a cluster divide synchronously. In real clusters, asynchronous cell divisions in a cluster can influence the cluster size distribution. To test whether asynchronous cell division significantly affects the predicted cluster size distribution, we simulated asynchronous cell division in the dynamic network. During a given division event, only a fraction of cells, chosen stochastically (based on the probability of cell division), were allowed to divide (**Figure S6I**). The probability of cell division per generation varied from 0.1 to 1. It only significantly affected cluster size distribution when the probability was less than 0.7. A probability value of 0.7 or less implies that a significant fraction (>30%) of the cells are lagging by unit division time which is more than the variation in doubling time for cells grown under the conditions of our experiments.

We tested the effect of varying the link breaking rate on cluster size distribution by using a strain whose chitin hydrolase (Cts1) expression can be tuned by altering the β-estradiol concentration in the media (**Figure 2G and S6D**). If links are hydrolyzed progressively, their fracture probability will increase with age, making older links more likely to break than younger ones. If clusters must break no later than after κ divisions, the cluster size distribution will follow the Gamma distribution; if the breaking rate (δ) is a contant, the probability that a linkage has survuived falls exponentially with its age. Because the Gamma distribution includes zero, which is not a cluster size, we used a zero-corrected version of the Gamma distribution: Erlang’s distribution. For random variables that follow Erlang’s distribution, the mean and standard deviation are linearly correlated, for our model, the slope should be 1/√*κ*. We observed that the means and the standard deviations of the cluster size distributions, produced at different levels of *CTS1* expression, followed Erlang’s distribution (**Figure 2H**), confirming the choice of exponential distribution for modeling the time for which a link survives.

Because the SA model predicts that clusters preferentially fragment at their older links, it implies that the two daughter clusters should be of similar size. The ratio between the size of the smaller of two daughter clusters (S_daughter_) and their mother cluster (S_mother_) gives an indication of whether the breaking mechanism is biased towards the oldest link. If the cluster always breaks at its oldest link the ratio is very close to 0.5. The SA model predicted that the distribution would be biased towards higher ratios. We tested this prediction by time-lapse imaging of *BUD4*-OFF clusters labeled with CFP (cyan fluorescent protein) during their growth and fragmentation cycles. (**Figure 3A**). The clusters were imaged at a low magnification to increase the depth of field, allowing us to use the total signal from a cluster as a proxy for the number of cells it contains (**Figure 3B)**. Consistent with our prediction, we found that the ratio (S_mother_/ S_daughter_) distribution is biased towards higher ratios (**Figure 3C**). We found an agreement when we performed similar estimations using the simulated dynamic network model (**Figure 3D, S7C and S7D**), increasing our confidence in the SA model as a phenomenological description of cluster growth and breakage.

**Figure 3.**
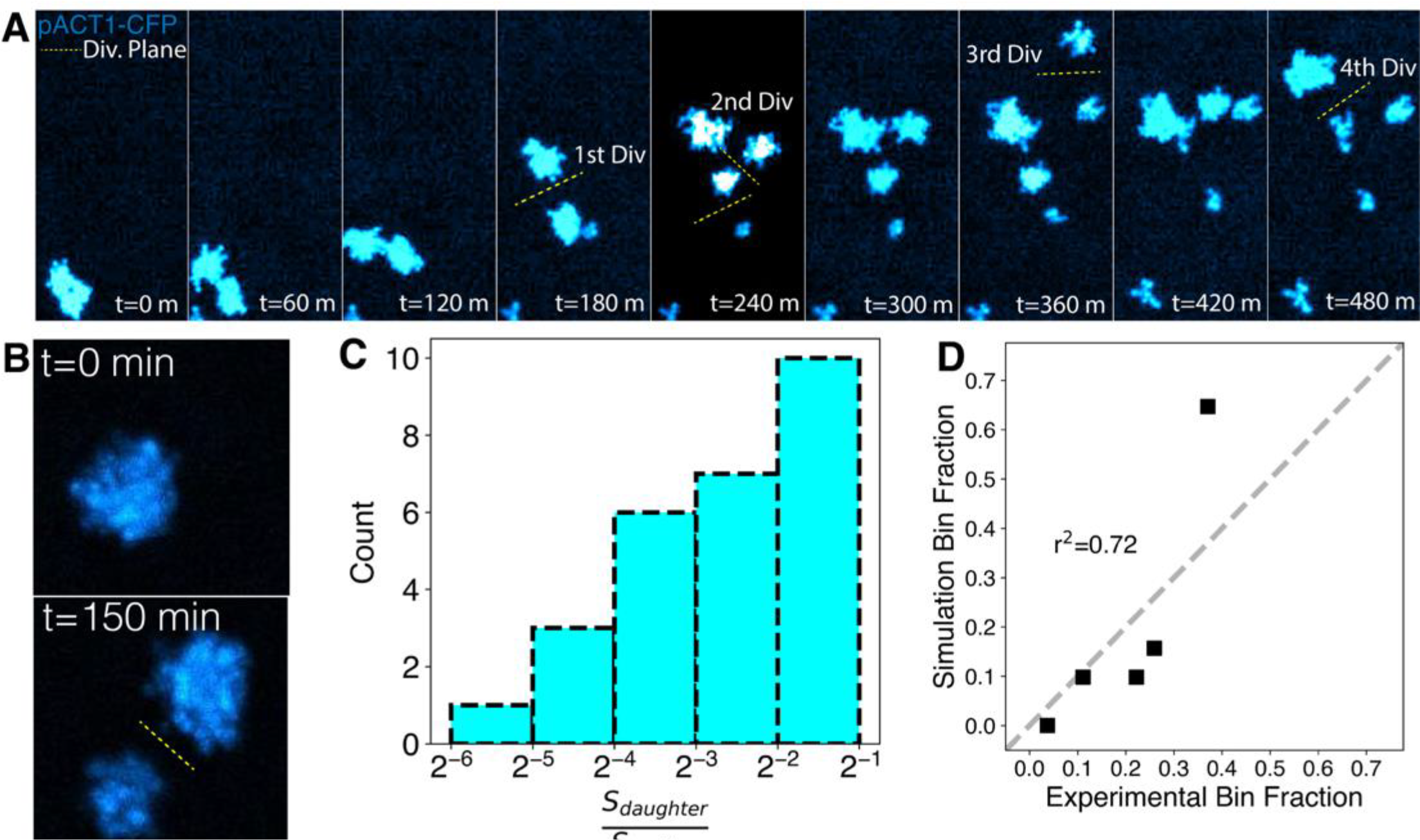
Tracking cluster growth and fracture shows agreement with SA model. A) Tracking lineages of clusters during growth-fragmentation cycles using a BUD4-OFF strain constitutively expressing CFP. The total fluorescence from the clusters is a measure of their size. B) The number of cells per cluster is estimated using their total fluorescence and cluster divisions are detected by the separation of clusters C) Distribution of the ratio of the mean number of cells per cluster in the mother to the smaller daughter cluster (n=28 mother-daughter pairs). D) Comparison of experimental and simulated mother: daughter cluster size ratios. Bin fraction is the fraction of data points in the specific bins defined in C.

### The ratio between growth rate and link breakage rate determines cluster size

We examined the effect of growth and link breaking rates on cluster size. We hypothesized that for a given kissing number and division rate, there is a threshold for the breaking rate: when the rate is below the threshold, the kissing number is the primary determinant of cluster size and clusters are big, when the rate is higher, links break before the number of connections reaches the kissing number and clusters are smaller (**Figure 4A**). The breaking rate threshold gets smaller as the division rate falls because links must persist longer to allow the clusters to reach their maximum possible size. We lowered the glucose concentration (to 0.02% or 0.05%) to reduce the mean cell division rate (**Figure S8B**). In addition, we uncoupled Cts1p production from its normal transcription factor, Ace2p^23,24^, in two ways: deleting *ACE2* or expressing *CTS1* from a β-estradiol regulated promoter. In both manipulations, the rate of chitin hydrolysis should be independent of the growth rate. We measured the cluster size distribution in a *BUD4*-OFF *ace2Δ* strain growing exponentially in different concentrations of glucose. Consistent with our hypothesis, the cluster size distribution shifted to bigger clusters (reflected in the mean of the distribution) in higher glucose concentrations (2% and 0.2%) (**Figure 4B**).

**Figure 4.**
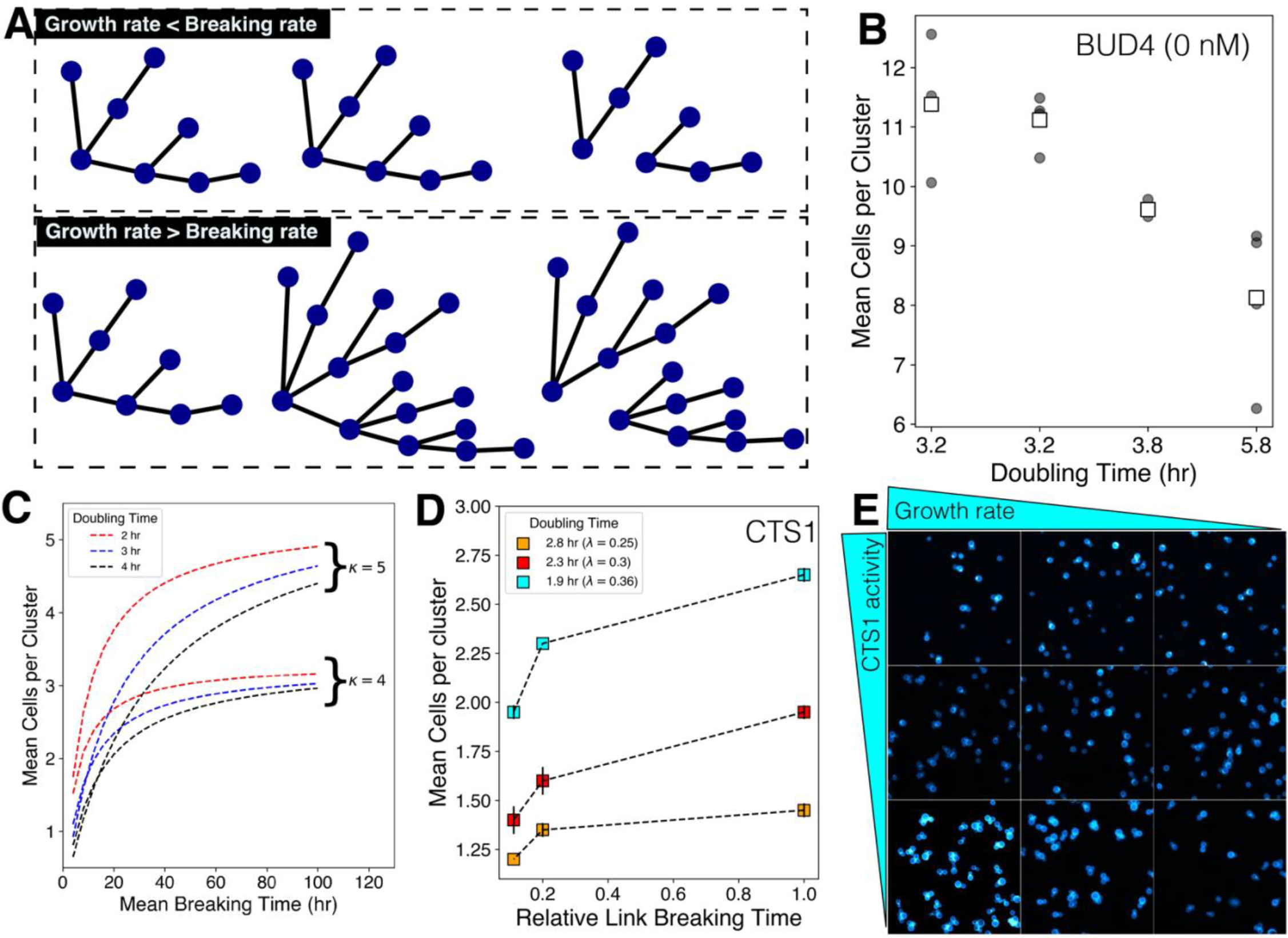
The relative values of cell division rate and link breaking rate sets the size of clusters. A) The hypothesis that a cellular growth rate higher than the link breaking rate yields clusters with a higher number of cells per cluster. B) Clusters with uninduced Bud4p (0 nM β-estradiol in a strain with *BUD4* expressed from a β- estradiol inducible promoter) show an increase in mean cluster size with an increase in cell division rate. Closed circles represent biological replicates and squares represent their means {replace BUD4 (0 nM) with “uninduced *BUD4*” in the figure} C) Simulations support a model in which the growth rate sets a threshold beyond which the breaking time cannot affect the mean cluster sizes. For two different kissing numbers, the mean cluster sizes reach a maximum value independent of cell division rate or link breaking rate. D) Experimental data for change in the mean number of cells per cluster with increasing growth rate and chitinase (Cts1p) activity. The mean size is calculated from two independent replicates E) Images of clusters for varying growth rates and chitinase induction.

To study the effect of growth rate on the mean cluster size, we explored parameter space and asked how the kissing number and link breaking rates influence the relation between growth rate and mean cluster size. Simulations suggested that, for a fixed cell division rate, the mean number of cells per cluster increases monotonically as the link breaking rate falls (**Figure 4C**). The mean number of cells per cluster, at half the value set by purely sterically induced breakage (set by the kissing number), is proportional to the doubling time of cells in the cluster (**Figure 4C)**. Thus the relative values of the link breaking rate (*δ*) and cell division rate (*λ*) alter the mean size of the clusters. In contrast, the kissing number (*κ*) sets the maximum value.

We simultaneously varied the link breaking rate (*δ*, by altering the estradiol concentration) and the cell division rate (*λ*, by varying glucose concentrations) to test the validity of this model. The cell division rate (*λ*) was calculated from the growth curves for bulk cultures in different glucose concentrations (**Figure S8C**). We examined all the combinations of three different ß-estradiol concentrations, 0, 4, and 8 nM, with three different cell division rates (*λ*), 0.22 hr^-1^, 0.18 hr^-1^, 0.12 hr^-1^. The mean number of cells per cluster was used as a proxy for the effect of cell division rates and link-breaking rate on cluster size distribution. At a given link breaking rate, increasing the growth rate increased cluster size, and at a given growth rate, increasing link breaking rate decreased cluster size (**Figure 4D and 4E**). We predicted that the effect of increasing the link breaking rate should be stronger at lower growth rates, but this is not what we see. Our interpretation is that there are two components to the link breaking rate, one due to the expression of *CTS1* and the other due to the expression of other hydrolytic enzymes. At the lowest growth rate, 0.12 hr^-1^, we argue that the *CTS1*-independent link breaking rate is already high enough to substantially diminish both the size of the clusters and the response to graded *CTS1* expression (**Figures 4D and 4E**).

### Higher kissing number minimizes compositional heterogeneity in differentiating multicellular clusters

A three-step model for the evolution of clonal multicellularity posits that the first step is the failure of cell separation, the second is the division of labor between cells, and the third is the separation of germ and somatic lineages, with irreversible differentiation from the faster dividing germ cells to more slowly dividing somatic cells. When the presence of both cell types maximizes the rate of reproduction, clusters that are all somatic cells are disadvantaged because they can never produce germ cells.

We studied how the parameters governing cluster size affect the distribution of germ and somatic cells using an engineered yeast strain that mimics irreversible differentiation^12^. In this strain, the germ cells express a cycloheximide resistance gene that can be used to set the relative division rates of germ and somatic cells, and Cre-mediated recombination converts germ cells into somatic cells by excising this gene and expressing invertase (Suc2p), which hydrolyzes sucrose extracellularly^12^, releasing glucose and fructose as a public good. The two cell states are distinguished by the fluorescent protein they express: germ cells express RFP and somatic cells express YFP. We define the switching rate from the germ state to the somatic state as *π*_*g*_→_*s*_and the relative growth advantage of germ cells with respect to somatic cells as *μ*_*r*_; *π*_*g*_→_*s*_can be experimentally modulated by changing the expression of Cre-recombinase using ß-estradiol and *μ*_*r*_ can be controlled by changing the cycloheximide concentration.

Sterile clusters, which contain only germ or only somatic cells, are formed when the link between a cell that has differentiated and its undifferentiated mother cell breaks (**Figure 5A**). When the kissing number is low, the number of cells connected to a given cell will be low. We therefore expect the probability of the differentiating and the fracturing branch coinciding to be high even if differentiation and breakage are independent processes. As the kissing number rises, this probability falls as there will be more branches to distribute the events among. We modified the SA model to incorporate the dynamics of differentiation. Briefly, when a new node is added to a growing network, it is assigned a phenotypic state (germ or somatic) based on the value of *π*_*g*_→_*s*_. Because differentiation is irreversible, *π*_*s*_→_*g*_=0. When germ cells have no growth advantage over somatic cells (*μ*_*r*_ =1), increasing the kissing number only marginally reduced the fraction of sterile clusters that solely contained germ cells (**Figure 5B**). When germ cells divide faster (*μ*_*r*_ > 1), the effect of kissing number on the fraction of sterile clusters was more pronounced (**Figure 5B**). Simulations also predicted that increased kissing number would increase the fraction of clusters having both cell types (mixed).

**Figure 5.**
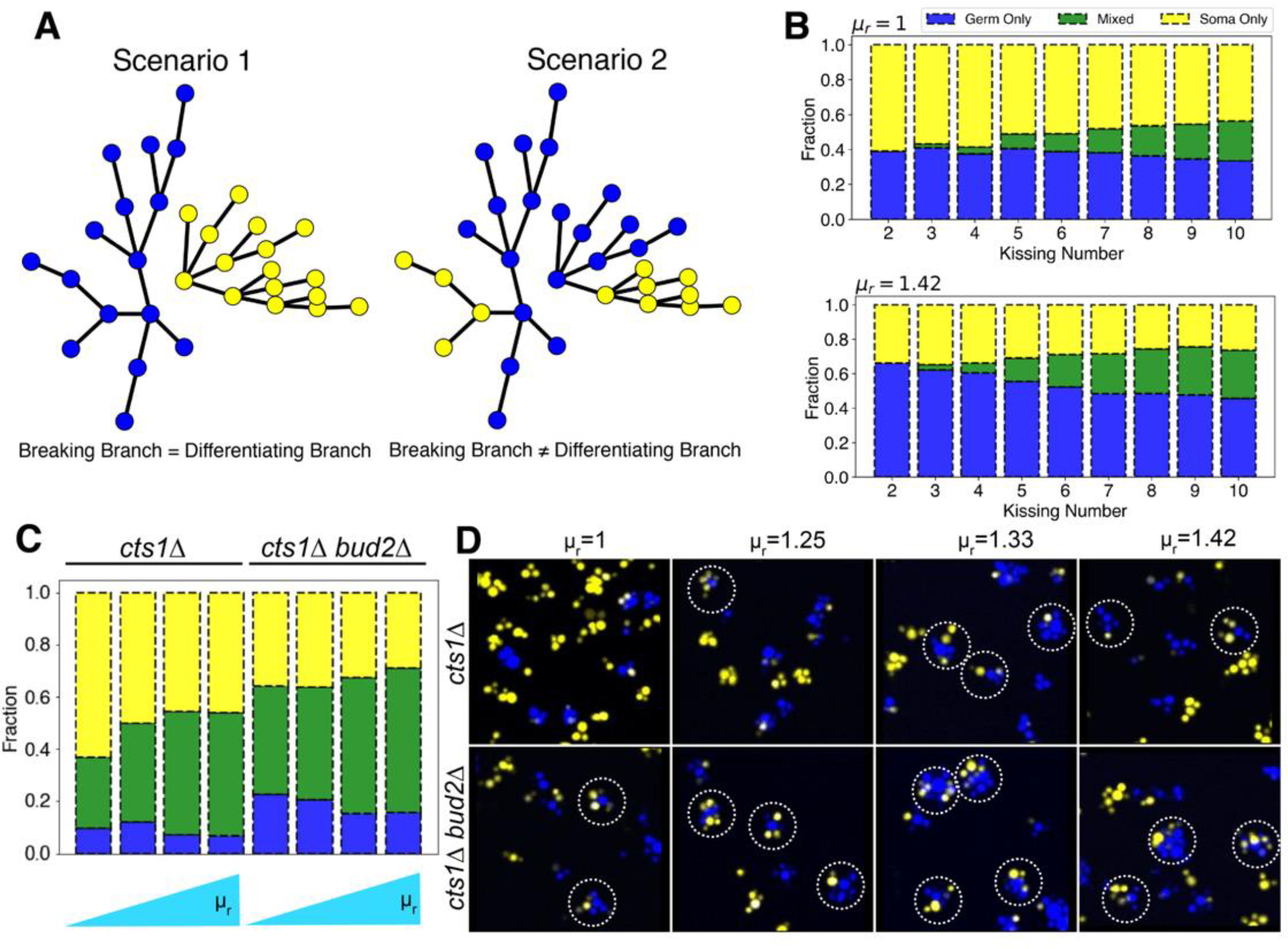
Kissing number (*κ*) affects compositional heterogeneity in differentiating clusters. A) Two scenarios for the fragmentation of clusters where cells can differentiate irreversibly: in scenario 1, one daughter cluster contains only germ cells and the other only somatic cells; in scenario 2, both daughter clusters contain both cell types. B) Simulations with the SA-based model. The effect of increasing the kissing number and relative growth advantage on proportion of cluster types: only germ, only somatic or mixed C) Experimental composition of clusters (germ only, somatic only, and mixed) with varied kissing number and relative growth rates determined experimentally. D) Images for clusters with two cell types for various conditions shown in (C); dotted circles show mixed clusters.

To experimentally test these hypotheses, we modified the kissing number of differentiating clusters. We could not delete *BUD4* because its effect on kissing number is dependent on the absence of *ACE2*^27^and *ACE2* is required for the differentiation system used here. Instead, we deleted either *BUD2*^25^ (*bud2*Δ cells bud at random sites on cell surface) or *BUD3*^26^ (*bud3*Δ cells bud at bipolar sites) in a *cts1*Δ background. These clusters have increased kissing numbers, and thus increased maximum attainable size, due to switch from axial budding to bipolar or random budding. The fraction of clusters that have exclusively germ or somatic cells or a mixture of both can be identified by flow cytometry based on the total RFP or YFP signal emitted for each cluster (**Figure S9**). We measured the fraction of each cluster type in glucose-containing medium at increasing concentrations of cycloheximide (which elevates *μ*_*r*_) for both *cts1*Δ, and *bud2*Δ *cts1*Δ clusters. Increasing the kissing number (comparing *cts1*Δ and *bud2*Δ *cts1*Δ clusters) reduced the fraction of sterile clusters (**Figure 5C**). It also led to a significant increase in fraction of mixed clusters. We confirmed these results using microscopy (**Figure 5D**). The fraction of clusters that are exclusively germ cells disagreed between the simulations and the experiments: while the model predicted that this fraction would fall at high kissing numbers, the experiment revealed an increase.

We asked whether the model parameters influence the ratio of the cell types in individual clusters, the variability in this ratio across clusters in the population, and whether they can be modulated in a predictable manner. In conditions where metabolic division of labor or cross-feeding occurs, the ratio of two cell-types in a cluster is particularly relevant. An unbalanced ratio would underfeed one of the two cell-types and reduce the performance of the cluster.

We modified our imaging pipeline (**Figure S2**) to measure the ratio of germ and somatic cells in each cluster. This change allowed us to determine the fraction of a given-cell type in individual clusters based on YFP and RFP signals across hundreds of clusters in a population and for several different perturbations. First, we tested how parameters specific to differentiating clusters i.e., *π*_*g*_→_*s*_and *μ*_*r*_alter this mean fraction of somatic cells per cluster. We express the net effect of these two parameters as *σ*_*g*_→_*s*_= *π*_*g*_→_*s*_/ *μ*_*r*_. Therefore, *σ*_*g*_→_*s*_, the net switching rate, describes the net effect of the formation (*π*_*g*_→_*s*_) and slower proliferation (*μ*_*r*_) of the somatic cell-type. The SA dynamic network model predicted that the mean fraction of somatic cell-type in clusters would monotonically increase with the net switching rate (**Figure 6A**). As the model predicts, any desired ratio could be attained by fixing the system at a given net switching rate (*σ*_*g*_→_*s*_). This relation seemed to be largely independent of both the kissing number (*κ*) and the link breaking rate (*δ*). To test the validity of this relation, we set-up an experimental system to alter the net switching rate by varying the concentrations of ß-estradiol and cycloheximide. We measured the fraction of somatic cells for hundreds of clusters per sample, for all combination of strains, switching rate and relative growth advantage, a total of 9 conditions. Consistent with model predictions, the mean somatic fraction across clusters scaled monotonically with net switching rate independent of the strain background (**Figure 6B**). The fraction of somatic cells changed from 0.25 for the lowest *σ*_*g*_→_*s*_to 0.6 for the highest *σ*_*g*_→_*s*_. We also explored the cluster-to-cluster variability for each condition. As this fraction is bounded between 0 and 1, we used the Gini coefficient to measure cluster-to-cluster variability in the fraction of somatic cells. The Gini coefficient is adopted from econometrics where it is used to quantify income variability within populations^28^. We found a strong negative correlation between mean somatic fraction (**Figure 6C**) and the Gini coefficient for all the conditions measured (r^2^=0.87). This implies a scaling relation between mean and variability like that observed for the mean and noise in gene expression^29^. A higher somatic fraction automatically leads to a reduction in cluster-to-cluster variability. We obtained a similar relationship using conventional measures like coefficient of variation (CV), which is typically used for unbounded measurements (**Figure S10B**).

**Figure 6.**
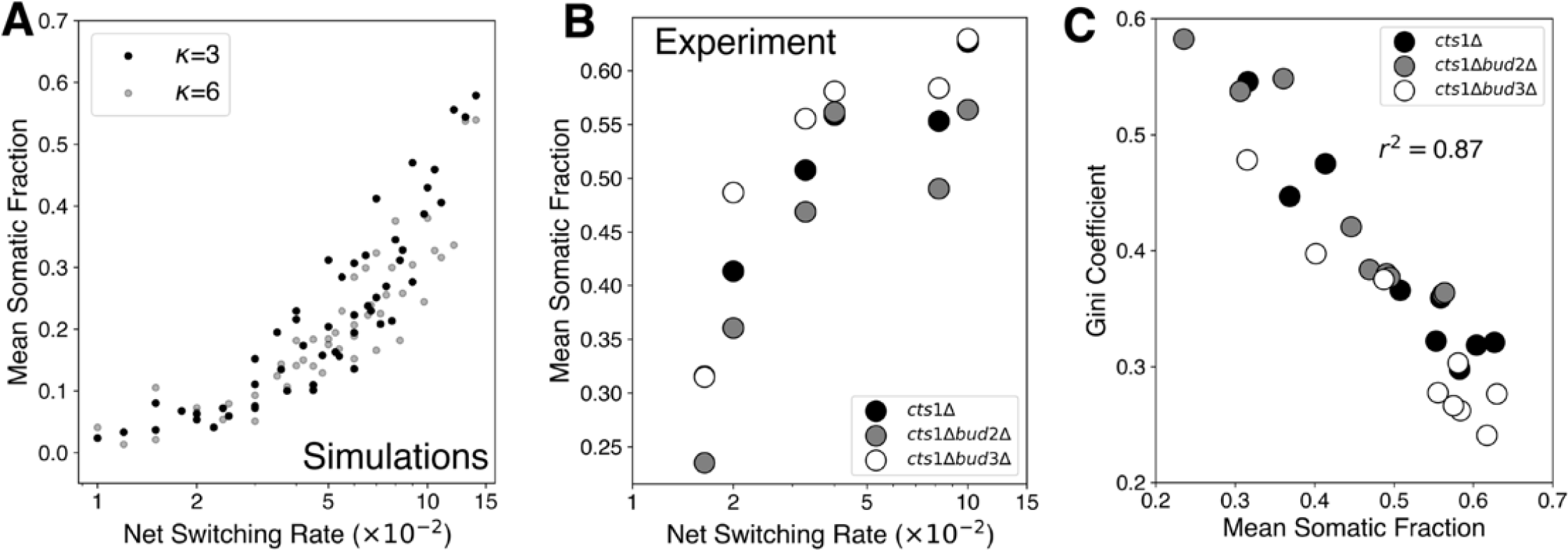
Average somatic fraction is negatively correlated with cluster-to-cluster compositional heterogeneity. A) Model predictions of mean somatic fraction as a function of net switching rate (*σ*_*g*_→_*s*_= *π*_*g*_→_*s*_/*μ*_*r*_) for two different kissing numbers B) Experimentally measured mean somatic fraction across three different genetic backgrounds and variable net switching rate. The net switching rate was controlled by cycloheximide and β-estradiol concentrations. C) The relationship between mean somatic fraction across clusters and cluster-to-cluster variability estimated by the Gini coefficient.

## Discussion

Multicellularity has evolved multiple times in nature and has been studied in the lab by experimental evolution and engineering. These studies prompted us to examine the variables that determine the size and composition of multicellular clusters in budding yeast using simulation and experiments. The model that best predicted the experimental data relied on a combination of steric constraints and an age-dependent fall in the probability of a link surviving. We found that clusters exist in two regimes: in one the link breaking rate is lower than the cell division rate and the kissing number sets the cluster size, whereas in the other, the link breaking rate is greater than the division rate leading to smaller clusters. Finally, the compositional heterogeneity of differentiating clusters was maximized by increasing the kissing number and the growth advantage of germ over somatic cells.

Experiments that have evolved multicellularity show remarkable differences in cluster size distributions across independent lines^10^. While several populations show convergent evolution, like the inactivation of *ACE2*, cluster size distribution varies across evolution lines^10^. Our simulations and experiments show that cluster size distribution and composition can be predicted from a small number of parameters. Multicellular growth cycles involve the growth of cells as an outcome of cellular metabolism and fractures of the linkage between of cells as an outcome of chemical decay of the links and physical forces. While the details of the location and magnitude of physical forces is an interesting question, it is dispensable for predicting cluster size distribution. We argue that the few phenomenological parameters used in our model, which can be directly linked to specific proteins or sets of proteins, are sufficient to recapitulate the experimentally observed distribution across a set of conditions. The model, with additional features, could potentially help in predicting which specific parameter (Kissing number, link breaking rate, or cell division rate) would be most likely to change to produce a cluster size distribution that maximizes fitness.

In this study, the dynamic network model was programmed such that all the three parameters, kissing number, link breaking rate, and cellular growth rate involved are independent of each other. One possible extension of this model is a case where the model parameters are interdependent. For example, using this model, one can simulate what would happen if the cell division ceased at the core of the cluster where high cell wall stress is present. In this scenario, the cell division rate (*λ*) is set to zero when the number of connections a cell makes reaches the kissing number (*κ*). Simulations suggest this relationship allows the clusters in the population to accommodate a greater number of cells (**Figure 7**) than would be possible if all the parameters were independent (SA model). We speculate that such correlations might be present in evolved multicellular clusters^10,30^ with sizes much greater than those observed in this study. The effect of increasing kissing number due to changes in cell aspect ratio must also be considered for testing this prediction. We did not model the effect of changes in cell aspect ratio on cluster size distribution due its non-linear relation to kissing number and the necessity to account for the physical location of cells within a growing cluster, two parameters which are difficult to experimentally measure across a population of clusters.

**Figure 7.**
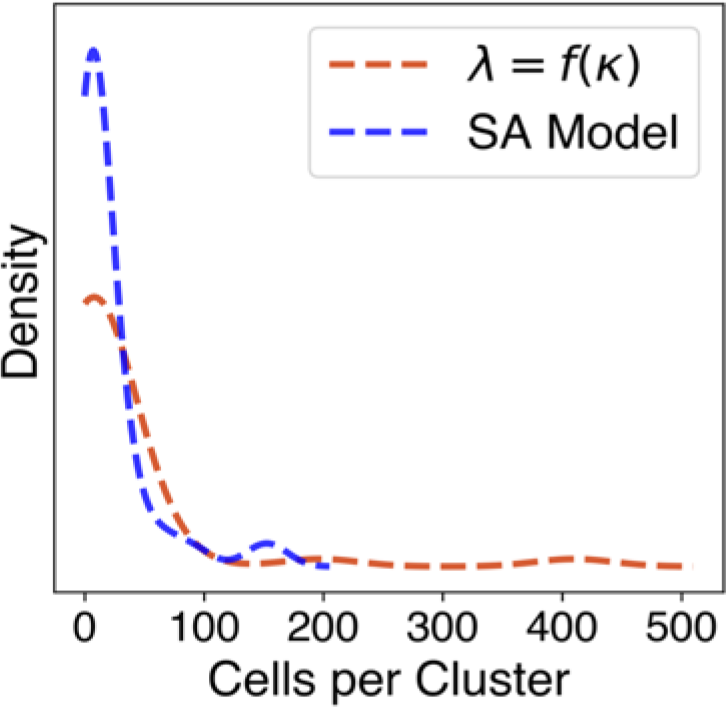
The model predicts kissing number dependent cell division rate allows clusters to accommodate greater number of cells. Simulated cluster size distribution from SA model and modified version where cells closer to reaching the kissing number stop dividing.

A limitation of our study is the assumption of tree-like growth of the clusters of cells which is idiosyncratic to multicellular yeast. In more complex life forms, the cells in the clusters are cemented together by an extracellular matrix comprising collagen, cellulose, and other polymers^31^. This allows cells to form connections with multiple other surrounding cells, unlike yeast clusters where a cell is only linked to its mother and its daughters. Moreover, cells in complex multicellular organisms can sense mechanical forces^32^ around them and use this information to control cell division. This regulation and external cues that control the rate and planes of cell divisions allows them to form non-symmetric 3D structures like organs. Given the ease of adding more features to the dynamic network model i.e., as attributes to nodes and edges, it would be interesting to ask what other features need to be incorporated to explain more sophisticated multicellular phenotypes. For example, would incorporating information about lateral inhibition^33^ as a node attribute explain how mechanosensitive circuits modulate multicellular cluster size when a high level of precision in required? Or would incorporating morphogen sensing^34^ as a node attribute based on geometric location allow us to predict cellular patterns?

## STAR Methods

### Key Resource Table

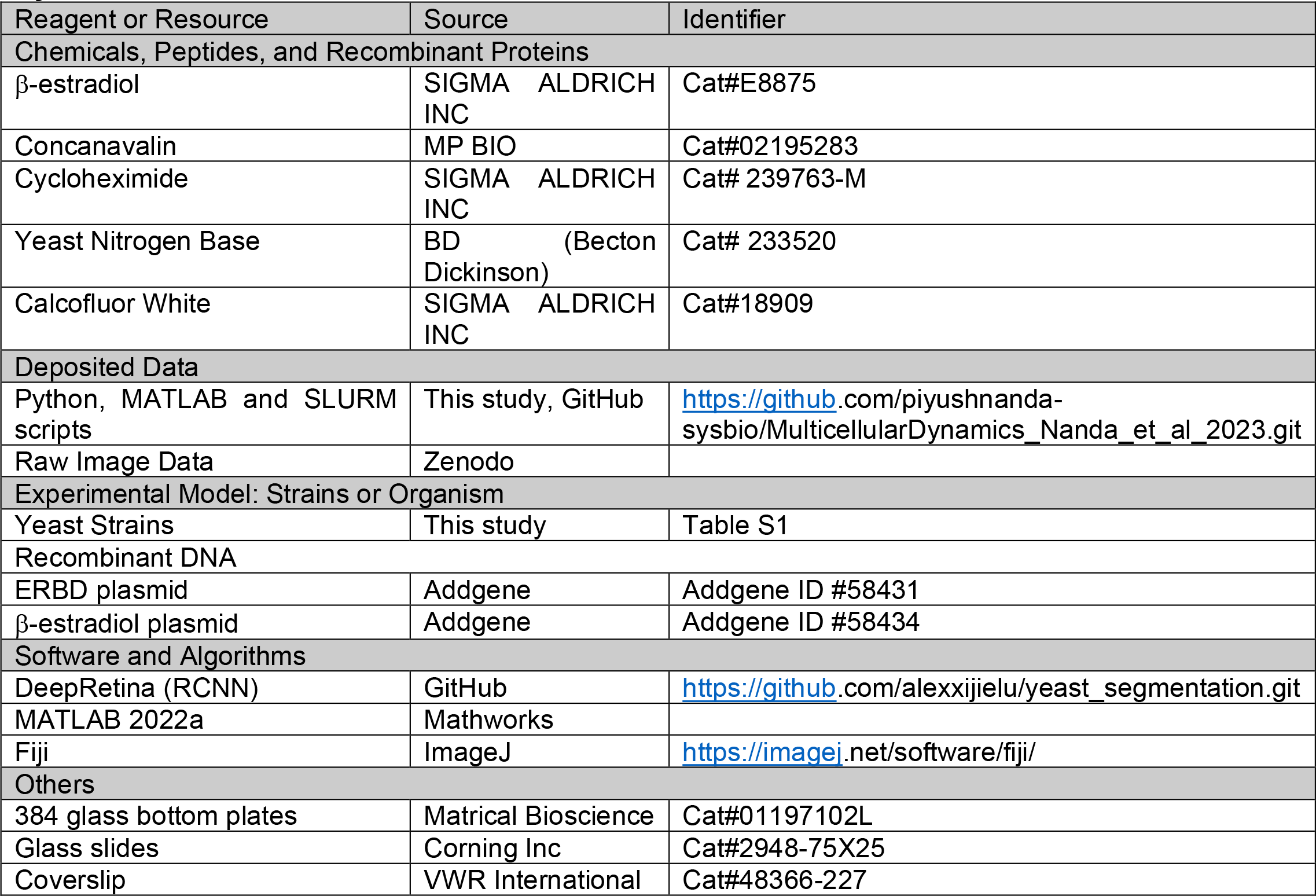

### Resource Availability

Further information or request for resources should be forwarded to the lead contact, Andrew Murray

### Materials Availability

Yeast strains and plasmid built for this project are available on request. Please contact the lead contact for further details.

### Data Availability

All scripts used for processing data or generating figures are hosted on GitHub. All raw data from microscopy experiments have been submitted in Zenodo. Any inquiry must be directed to the lead contact.

### Experimental Model and Subject Details

#### Yeast strains and media

All experiments were performed using strains constructed from a modified W303 background^35^. Briefly, the functional version of *BUD4* allele from S288C was engineered into the endogenous locus in W303. The β- estradiol-responsive transcription factor was introduced at the *HIS3* locus through an integrative plasmid^22^. Genes of interest were put under the control of β-estradiol promoter by replacing their endogenous promoter by homologous recombination. Constitutively expressed fluorescent markers were introduced at the *HO* locus using homologous recombination through the KanMX marker^36^.

Gene deletions were performed by amplifying the KanMX resistance marker with 500 bp upstream and downstream sequence from yeast deletion collection^37,38^ and introduced in the background of interest by homologous recombination. Confirmation PCR was performed to both check the integration of the resistance cassette and removal of target gene. Replica plating was performed for each successive transformation to confirm retention of previous markers.

Strains were grown in synthetic minimal media supplemented with the specified carbon source at various concentrations. Media were prepared freshly by diluting 10X stocks of the refrigerated constituents and prewarmed before inoculation. For every experiment, strains were freshly streaked out from −70°C glycerol stocks (15%) and allowed to recover for at least 2 days at 30°C. Strains were cultured in desired media for at least 10 generations before any measurements were performed. For some specific experiments, strains were grown for longer than 10 generations by maintaining them in exponential phase to allow the cluster size distribution to reach steady state. All the experiments were performed with two biological replicates for high-throughput imaging of size distributions. Replicate correlations have been described in supplementary figures.

### Methods details

#### Estimating number of cells per clusters through high-throughput imaging

Simulations using the models described in the main text produce a cluster size distribution, which we compared with experimental distributions. It is challenging to image individual cells in an entire three-dimensional cluster due to excessive light scattering and interference by the yeast cell wall. To overcome this challenge, the clusters were stained with calcofluor white (CW) and squeezed between a coverslip and a slide. The fluid volume between the coverslip and slide was optimized so that the cells from a single cluster formed a monolayer without lysing (**Figure S2**). We performed a timelapse of clusters disintegrating to a monolayer to check for any lysis event or mixing of cells from different clusters **(Video S1**). By optimizing the density of clusters in a field of view, we spatially resolved cells belonging to individual clusters in a monolayer (**Figure S2 and Figure S3**). We used a previously established Mask-RCNN pipeline (Region-based Convolutional Neural Network) images to segment yeast cells using images obtained from the CW channel (**Figure S2**). The codes and associated files were acquired from https://github.com/alexxijielu/yeast_segmentation. We collected multiple fields of view (>60 FOVs, i.e., ∼200 clusters) to obtain an accurate distribution of cluster sizes. The minimum number of clusters to be obtained to claim a detectable difference in two conditions or genetic backgrounds was calculated based on Cohen’s *d* (**Figure S2A**). For all the experiments, the expected effect size was <0.4 and therefore, 200 clusters were enough to derive a representative distribution. The centroid of cells in segmented images was estimated and used to cluster cells based on their spatial location using DBSCAN (Density-Based Spatial Clustering of Applications with Noise). The number of cells corresponding to each cluster was used to generate a discrete distribution of cluster sizes (**Figure S2**). We obtained reproducible cluster distributions for various genetic backgrounds (corresponding to size distributions) (**Figure S2B-D).**

Strains were grown in the indicated media for at least 10 generations to reach their steady state distribution of cluster sizes. For imaging, exponentially grown cells were incubated with 1 volume of calcofluor white for 1 minute at room temperature. Approximately, 2.8 uL of the mixture was squeezed between a microscope slide and a 20 mm × 20 mm coverslip. The coverslip and microscope slide were cleaned with an air jet before experiments to remove dust particles. A Nikon Eclipse TiE inverted fluorescence microscope with a Hamamatsu EMCCD camera was used for all imaging. The microscope was controlled through MetaMorph software. An automated script written in MetaMorph scanned 49 positions on the slides in an arrayed fashion. The density of cultures was optimized to yield at least 4-5 clusters per field of view. Monolayer formation was manually verified for each sample by imaging z-stacks and confirming the absence of more than one layer of cells.

#### Timelapse imaging

All timelapse imaging was performed using the microscope described above. Concanavalin-A coated wells in a 384 well plate were used to perform lineage tracing of cluster growth and division using a 10X objective with 10 mm working distance. Fluorescence from a constitutively expressed CFP was used as a proxy for total number of cells per cluster. The depth of field was calculated using excitation wavelength and magnification values. The plates were filled with at least 40 uL of media to reduce evaporative loss. Strains were grown in specified media for 10 generations and were spun down on a 384 well plate at 300 g for 2 mins. Images were taken every 10 mins for a 12 h period.

#### Confocal Laser Scanning Microscopy of 3D clusters

A Zeiss LSM900 microscope was used for 3D imaging of multicellular clusters. Briefly, cells were stained with calcofluor white and loaded onto a concavity slide in 0.2% agarose to keep clusters stationary during imaging. The clusters were imaged using the DAPI channel with a z-stack width of 0.2 um. The 3D images were rendered using Fiji (ImageJ) and custom scripts.

#### Flow cytometry and sorting

All flow cytometry experiments were performed using a BD LSR II. The photomultiplier tube (PMT) voltages for forward and side scatter were optimized for each experiment to record maximum signal from the biggest clusters. The area signal from forward scatter (FSC-A) was used as a proxy for cluster size for preliminary measurements. The flow cytometry tubes were vigorously vortexed to obtain a uniform suspension of clusters before loading the samples.

Cell sorting was performed on multicellular clusters using a BD FACS Aria II using signal from the FSC-A channel. The FSC-A distribution was segmented into 4 quartiles and sorted into 4 different collection tubes at RT with minimal media. The cluster fractions were grown in flasks at 30 deg C for 48 hours. The evolution of the size distribution over time was recorded every 12 hours using flow cytometry.

#### β-estradiol titration

Strains were grown overnight in minimal media without β-estradiol. They were sub-cultured next day into minimal media with desired beta-estradiol concentrations. β-estradiol stocks were prepared in 100% ethanol and stored at −20°C. The stock was diluted to desired concentrations in minimal media. All measurements were performed after overnight growth (at least 10 generations) in presence of inducer. The dynamic range of inducer concentrations was estimated by performing a log-scale dose-response curve with the cluster size distribution as the output.

#### β-estradiol Cre induction and differentiation

Strains were freshly streaked out from −70°C glycerol stock. Single colonies were picked and checked for spontaneous Cre recombination events. Colonies with the pure germ state were grown in complete synthetic media (prepared from 10X stocks of constituents) overnight and diluted 100-fold into fresh media supplemented with desired concentrations of β-estradiol (inducing Cre recombinase expression) and cycloheximide. Cycloheximide stocks were made in 100% ethanol and stored at −20°C. The strains were grown for at least 14 generations (two transfers) to allow clusters to reach steady state fraction and size. Differentiation was confirmed by microscopy and flow cytometry.

The ratio of cell types in individual clusters was determined using a modified version of method described in Figure 2. Instead of calcofluor white, a constitutively expressed CFP present in both germ and somatic was used as segmentation marker and RFP (germ) and YFP (somatic) were used to identify the cell types. The number of cells of a given cell-type was estimated using the pipeline described in the quantification and analysis section.

### Quantification and statistical analysis

#### Dynamic Network Simulations and Analysis

Custom scripts were written in Python to perform dynamic network simulations using the NetworkX library (https://github.com/networkx/networkx). Briefly, a recursive program was run in which nodes (cells) were added every division time to extant nodes. Cell properties were encoded in node attributes and link properties were encoded in edge attributes. A function was used to update the attribute based on a trigger event: i) whether a kissing number was reached ii) the links reached a point of breaking. The network growth is initiated from a single cell but continues for 10 generations, after which at least 150 clusters have formed. Disconnected modules in the network were determined by finding set of nodes which don’t have connections with outside nodes. A disconnected module was identified as a cluster and the number of cells it contained was determined. Every simulation was performed for multiple random number seeds. The same random number seed was used for comparing effects of two different parameter sets on network properties.

#### Segmentation and estimation of cluster attributes

Segmentation of cells in clusters were performed using a yeast optimized pipeline based on DeepRetina. The pipeline was executed in a supercomputing cluster (FASRC) using custom SLURM scripts and images acquired were directly transferred for segmentation without any preprocessing or selection. The aspect ratio of images was preserved, and images acquired through the DAPI channel or CFP was used for segmentation.

MATLAB 2022B was used for image processing and analysis. Briefly, centroids corresponding to cells in a cluster was determined and used for performing DBSCAN (Density Based Spatial Clustering And Noise). DBSCAN assigns the cells to specific cluster based on its geometric location. Clusters with cells close to the image border (in the 50-pixel proximity) were rejected from downstream analysis pipeline.

For estimating the fraction of somatic cells per cluster, the signal from YFP channel was used. Briefly, the cutoff for YFP ON was determined by thresholding. Cells having YFP intensity greater than the threshold were assigned ‘somatic state’. DBSCAN was performed as described earlier and fraction of cells belonging to somatic category was determined. The signal from RFP channel (corresponding to germ state) was not used due to remnant signal from degraded protein that interfered with accurate determination of germ cell status.

The MATLAB script was executed from SLURM in a supercomputing cluster and all images from the same experiment were processed in the same batch. No manual filtering of images were performed at any stage of this pipeline.

#### Measuring mother-daughter cluster size correlation

The net fluorescence from individual clusters was determined using Fiji and lineages were manually assigned. A custom script written in Python was used to calculate the mother cluster-daughter cluster size correlations. The total size of mother cluster was estimated by summing up the sizes of daughter cluster right after the breaking event was recorded.

#### Estimating fraction of somatic, germ and mixed clusters

Flow cytometry files were analyzed using FlowCytometryTools library in Python3.7. The threshold for YFP and RFP channel was determined from the histogram of size (FSC-A) normalized fluorescence values. A cluster was assigned ‘Somatic only’ type if the YFP intensity was greater than the threshold and RFP intensity was less than the threshold. Similarly, a cluster was assigned ‘Mix’ type if both the intensity values were greater than the corresponding thresholds. Separate thresholds were determined for different samples and strains.

#### Statistical calculations

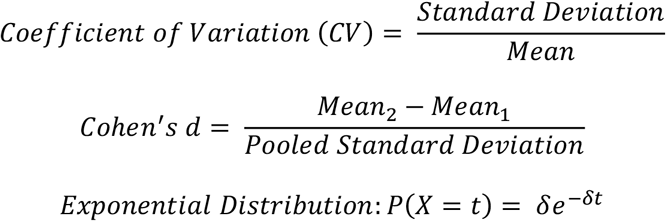

## Acknowledgements

The authors thank all members of the Murray lab for their feedback during lab meetings. We would also like to thank Boris I. Shraiman, Michael T. Laub, Jane Kondev, Allon Klein and Michael M. Desai for critically reading the manuscript and for their feedback. We want to thank the Harvard F.A.S. staff, especially Zach Niziolek, from the Bauer flow cytometry core facility, for assistance in FACS and flow cytometry assistance. We thank Christian Hellriegel from Harvard Biological Imaging Center (HCBI)/Zeiss. We sincerely thank Dr and Mrs Daniel Simmons for awarding Simmons Award to P.N. for performing advanced microscopy in HCBI. P.N., J.B. and A.W.M. were supported by the grant NIH/NIGMS R01 GM043987. A.W.M. is also supported by the NSF-Simons Center for the Mathematical and Statistical Analysis of Biology (N.S.F. #1764269, Simons #594596). T.L. was the Candy and William Raveis Fellow of the Damon Runyon Cancer Research Foundation while this research was performed.

## Reference

1 Grosberg RK, Strathmann RR. The evolution of multicellularity: A minor major transition? Annu Rev Ecol Evol Syst 2007; 38: 621–654.

2 Kirk DL. A twelve-step program for evolving multicellularity and a division of labor. BioEssays 2005; 27: 299–310.

3 Tong K, Bozdag GO, Ratcliff WC. Selective drivers of simple multicellularity. Curr Opin Microbiol 2022; 67: 102141.

4 Duran-Nebreda S, Solé R. Emergence of multicellularity in a model of cell growth, death and aggregation under size-dependent selection. J R Soc Interface 2015; 12. Doi:10.1098/rsif.2014.0982.

5 Knoll AH. The multiple origins of complex multicellularity. Annu Rev Earth Planet Sci 2011; 39: 217–239.

6 Brunet T, King N. The Origin of Animal Multicellularity and Cell Differentiation. Dev Cell 2017; 43: 124–140.

7 Staps M, van Gestel J, Tarnita CE. Emergence of diverse life cycles and life histories at the origin of multicellularity. Nat Ecol Evol 2019; 3: 1197–1205.

8 Koschwanez JH, Foster KR, Murray AW. Sucrose utilization in budding yeast as a model for the origin of undifferentiated multicellularity. PloS Biol 2011; 9. Doi:10.1371/journal.pbio.1001122.

9 Ratcliff WC, Denison RF, Borrello M, Travisano M. Experimental evolution of multicellularity. Proc Natl Acad Sci U S A 2012; 109: 1595–1600.

10 Koschwanez JH, Foster KR, Murray AW. Improved use of a public good selects for the evolution of undifferentiated multicellularity. Elife 2013; 2013. Doi:10.7554/eLife.00367.

11 Libby E, Ratcliff W, Travisano M, Kerr B. Geometry Shapes Evolution of Early Multicellularity. PloS Comput Biol 2014; 10. Doi:10.1371/journal.pcbi.1003803.

12 Wahl ME, Murray AW. Multicellularity makes somatic differentiation evolutionarily stable. Proc Natl Acad Sci U S A 2016; 113: 8362–8367.

13 Jacobeen S, Pentz JT, Graba EC, Brandys CG, Ratcliff WC, Yunker PJ. Cellular packing, mechanical stress and the evolution of multicellularity. Nat Phys 2018; 14: 286–290.

14 Márquez-Zacarías P, Pineau RM, Gomez M, Veliz-Cuba A, Murrugarra D, Ratcliff WC et al. Evolution of Cellular Differentiation: From Hypotheses to Models. Trends Ecol Evol 2021; 36: 49–60.

15 Bozdag GO, Libby E, Pineau R, Reinhard CT, Ratcliff WC. Oxygen suppression of macroscopic multicellularity. Nat Commun 2021; 12. Doi:10.1038/s41467-021-23104-0.

16 Jacobeen S, Graba EC, Brandys CG, Day TC, Ratcliff WC, Yunker PJ. Geometry, packing, and evolutionary paths to increased multicellular size. Phys Rev E 2018; 97: 1–6.

17 Mikhailov K V., Konstantinova A V., Nikitin MA, Troshin P V., Rusin LY, Lyubetsky VA et al. The origin of Metazoa: A transition from temporal to spatial cell differentiation. BioEssays 2009; 31: 758–768.

18 Pen I, Flatt T. Asymmetry, division of labour and the evolution of ageing in multicellular organisms. Philosophical Transactions of the Royal Society B: Biological Sciences 2021; 376. Doi:10.1098/rstb.2019.0729.

19 White MD, Zenker J, Bissiere S, Plachta N. How cells change shape and position in the early mammalian embryo. Curr Opin Cell Biol 2017; 44: 7–13.

20 Kang PJ, Hood-DeGrenier JK, Park HO. Coupling of septins to the axial landmark by bud4 in budding yeast. J Cell Sci 2013; 126: 1218–1226.

21 Ziv N, Siegal ML, Gresham D. Genetic and nongenetic determinants of cell growth variation assessed by high-throughput microscopy. Mol Biol Evol 2013; 30: 2568–2578.

22 Ottoz DSM, Rudolf F, Stelling J. Inducible, tightly regulated and growth condition-independent transcription factor in Saccharomyces cerevisiae. Nucleic Acids Res 2014; 42. Doi:10.1093/nar/gku616.

23 King L, Butler G. Ace2p, a regulator of CTS1 (chitinase) expression, affects pseudohyphal production in Saccharomyces cerevisiae. Curr Genet 1998; 34: 183–191.

24 Sbia M, Parnell EJ, Yu Y, Olsen AE, Kretschmann KL, Voth WP et al. Regulation of the yeast Ace2 transcription factor during the cell cycle. Journal of Biological Chemistry 2008; 283: 11135–11145.

25 Park HO, Chant J, Herskowitz I. BUD2 encodes a GTPase-activating protein for Budl/Rsrl necessary for proper bud-site selection in yeast. Nature 1993; 365: 269–274.

26 Chant J, Mischke M, Mitchell E, Herskowitz I, Pringle JR. Role of Bud3p in producing the axial budding pattern of yeast. Journal of Cell Biology 1995; 129: 767–778.

27 Voth WP, Olsen AE, Sbia M, Freedman KH, Stillman DJ. ACE2, CBK1, and BUD4 in budding and cell separation. Eukaryot Cell 2005; 4: 1018–1028.

28 Lerman RI. A Note on the Calculation the Gin1 Index. Econ Lett 1984; 15: 363–368.

29 Cai L, Friedman N, Xie XS. Stochastic protein expression in individual cells at the single molecule level. Nature 2006; 440: 358–362.

30 Bozdag GO, Zamani-Dahaj SA, Kahn PC, Day TC, Tong K, Balwani AH et al. De novo evolution of macroscopic multicellularity. bioRxiv 2021; : 2021.08.03.454982.

31 Dolega ME, Monnier S, Brunel B, Joanny JF, Recho P, Cappello G. Extra-cellular matrix in multicellular aggregates acts as a pressure sensor controlling cell proliferation and motility. Elife 2021; 10: 1–33.

32 Stassen OMJA, Ristori T, Sahlgren CM. Notch in mechanotransduction – from molecular mechanosensitivity to tissue mechanostasis. J Cell Sci 2021; 133. Doi:10.1242/jcs.250738.

33 Sancho R, Cremona CA, Behrens A. Stem cell and progenitor fate in the mammalian intestine: Notch and lateral inhibition in homeostasis and disease. EMBO Rep 2015; 16: 571–581.

34 Toda S, Mckeithan WL, Hakkinen TJ, Lopez P. Engineering synthetic morphogen systems that can program multicellular patterning. Science (1979) 2020; 331: 327–331.

35 Ralser M, Kuhl H, Ralser M, Werber M, Lehrach H, Breitenbach M et al. The Saccharomyces cerevisiae W303-K6001 cross-platform genome sequence: Insights into ancestry and physiology of a laboratory mutt. Open Biol 2012; 2. Doi:10.1098/rsob.120093.

36 Wach A, Brachat A, Pöhlmann R, Philippsen P. New heterologous modules for classical or PCR-based gene disruptions in Saccharomyces cerevisiae. Yeast 1994; 10: 1793–1808.

37 Winzeler EA, Shoemaker DD, Astromoff A, Liang H, Anderson K, Andre B et al. Functional characterization of the S. cerevisiae genome by gene deletion and parallel analysis. Science (1979) 1999; 285: 901–906.

38 Giaever G, Chu AM, Ni L, Connelly C, Riles L, Véronneau S et al. Functional profiling of the Saccharomyces cerevisiae genome. Nature 2002; 418: 387–391.

