## Supplementary Figures for "Multicellular growth as a dynamic network of cells"

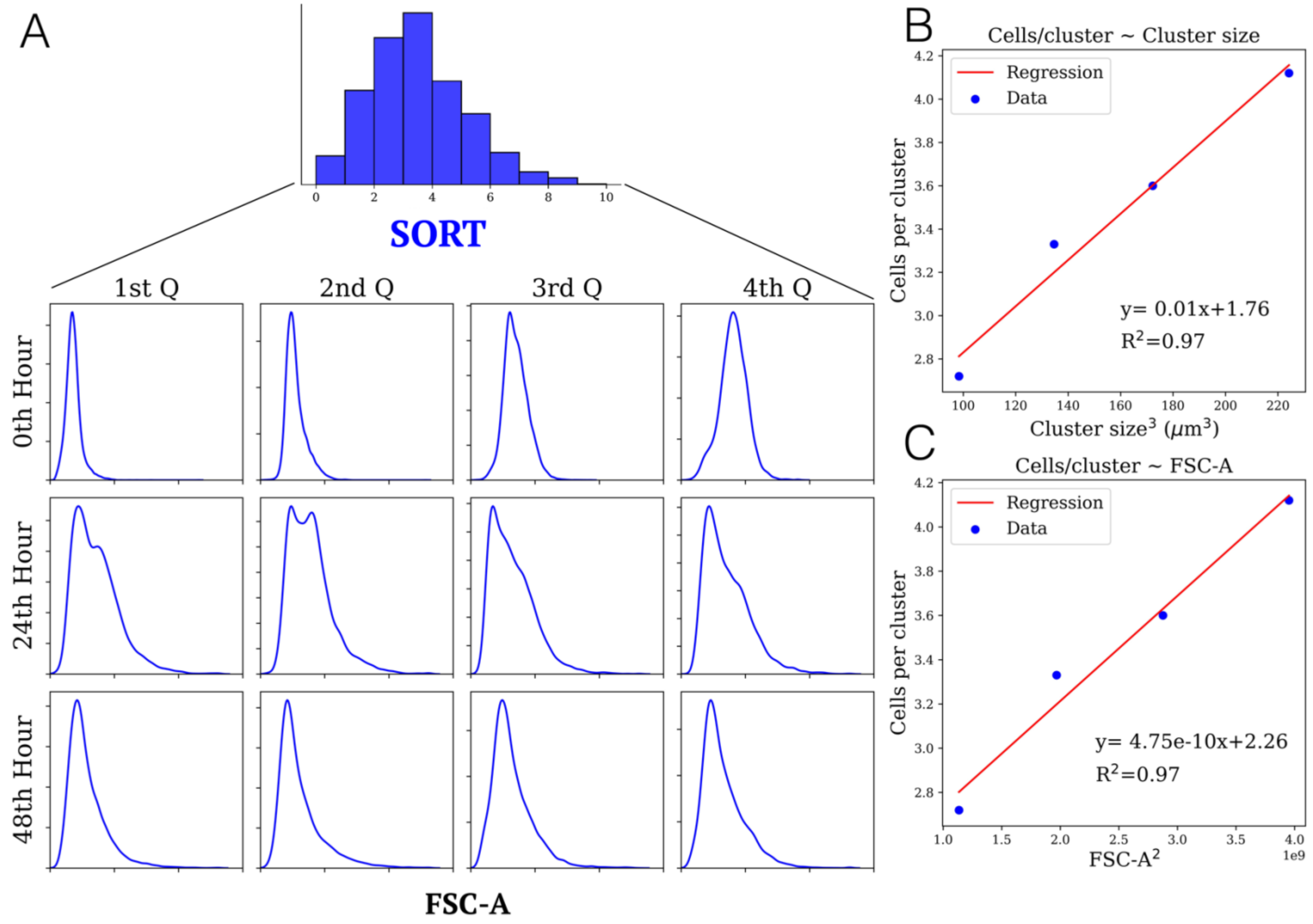


**Figure S1** A) Evolution of cluster size distribution over 48 h growth for four fractions (quartiles) sorted from clusters with reduced Cts1 expression. The x-axis shows the forward scatter area measured by flow cytometry  B) Correlation between mean cluster size measured by counting cells per clusters and multisizer based volume measurement. C) Correlation between mean cluster size measured by counting cells per clusters and flow cytometry.

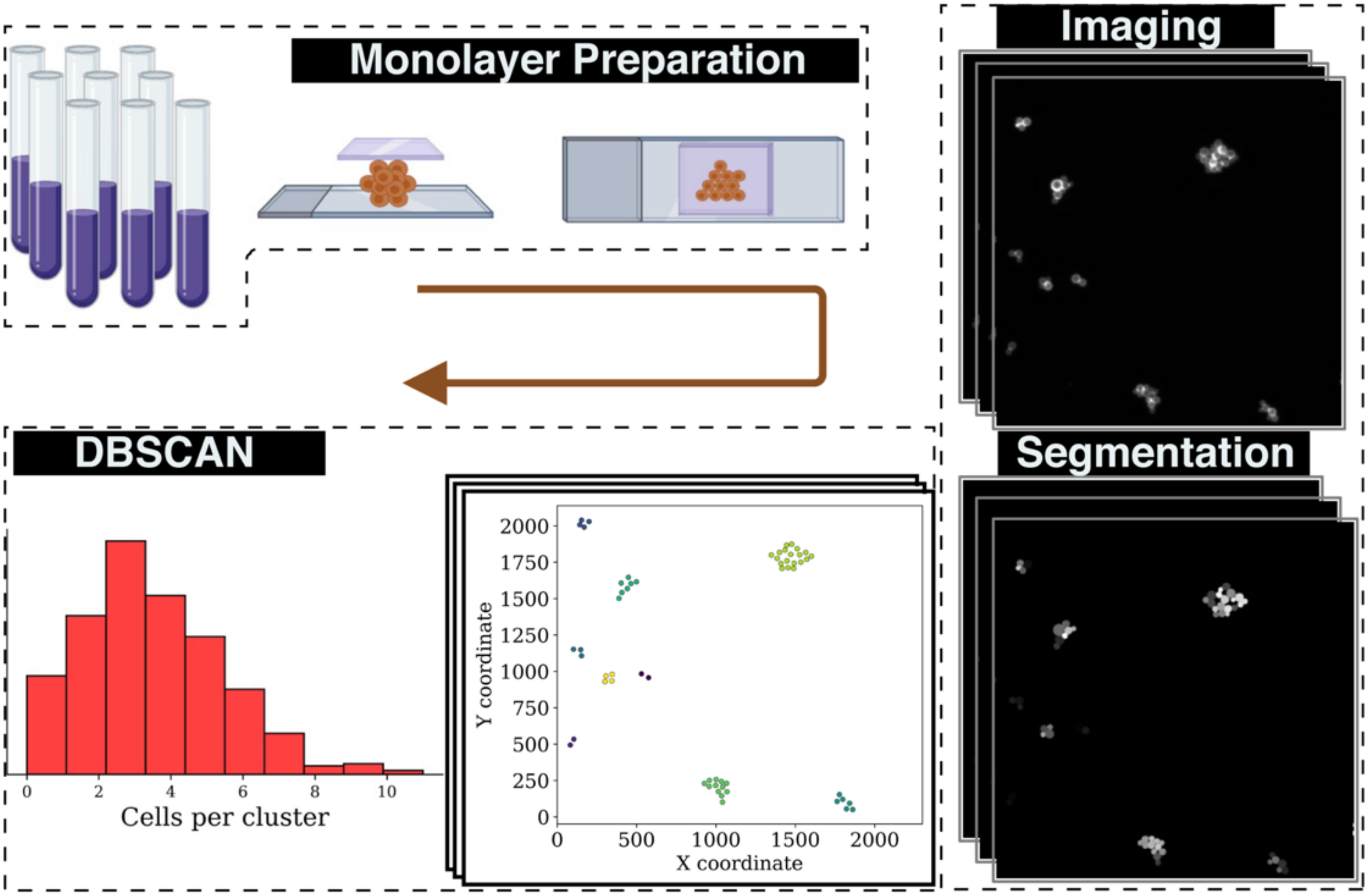


**Figure S2 A high-throughput pipeline to estimate the number of cells in each cluster** Clusters are grown in conditions of interest, squeezed between a coverslip and microscope slide, and imaged using light microscopy. The cells are segmented using an RCNN-based (Region based Convolutional Neural Network) segmentation pipeline. Centroid coordinates are used to identify clusters (using density-based spatial clustering of applications with noise (DBSCAN)) and the number of cells per group for hundreds of clusters is estimated.

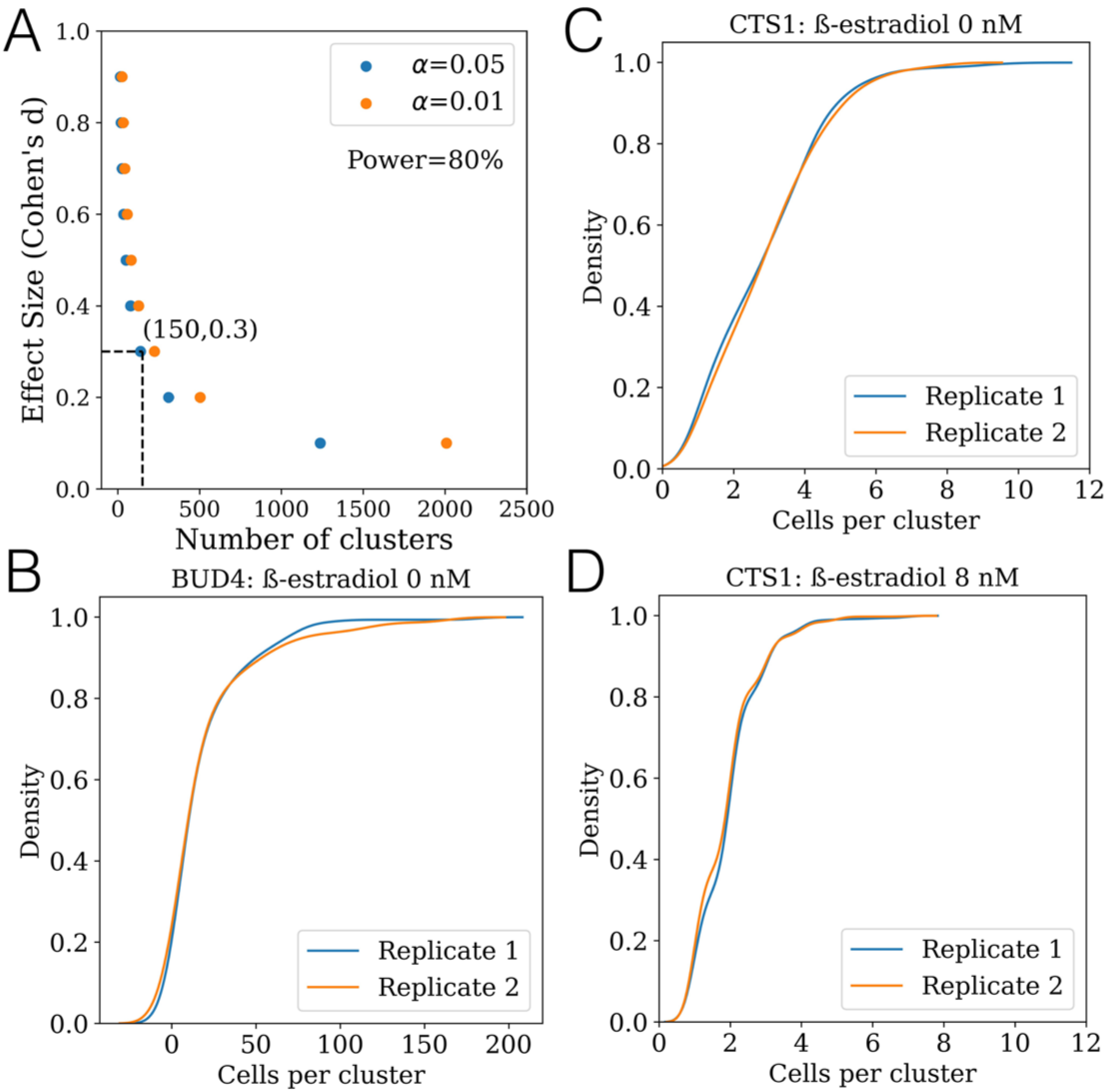


**Figure S3** A) Estimated number of clusters required to test for a certain effect size predicted by Cohen's d (the difference between two means divided by the standard deviation of the pooled data). B) Cumulative density function (CDF) for cluster size distributions for uninduced Bud4p in an *ace2*Δ strain, in which Bud4 is expressed from a β-estradiol-activated promoter, for two independent replicates C) CDF for cluster size distributions for uninduced Cts1p, in a strain in which Cts1 is expressed from a β-estradiol-activated promoter, for two independent replicates D) CDF for cluster size distributions for partially induced (8 nM β-estradiol) Cts1p for two independent replicates.

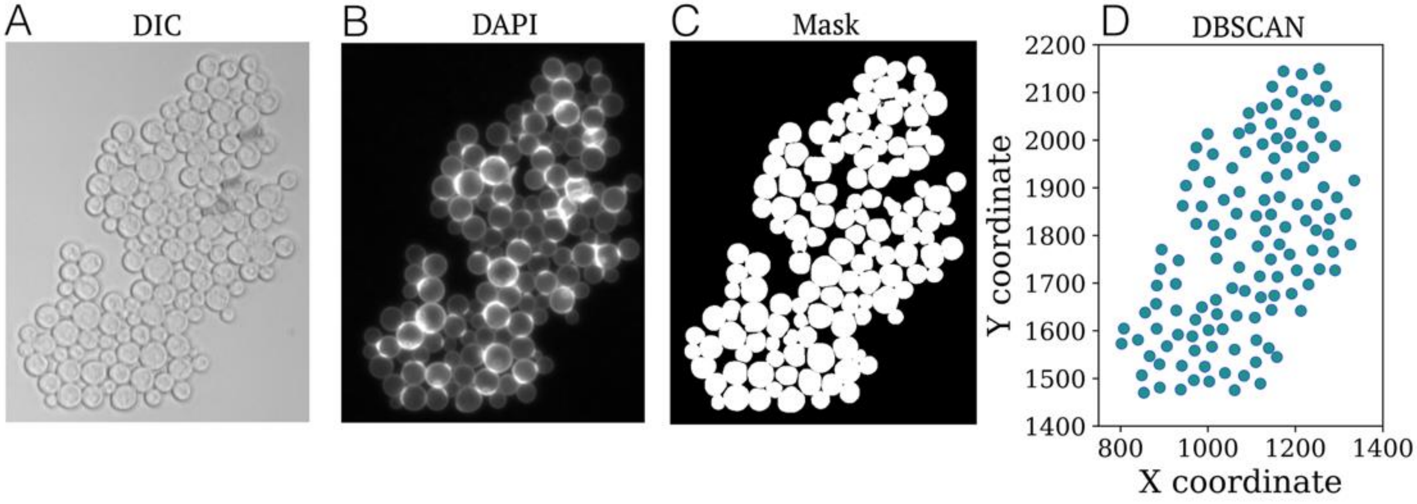


**Figure S4** Example images of a cluster from an *ace2*Δ strain with uninduced Bud4p, expressed from a β-estradiol-activated promoter A) DIC image B) DAPI C) Mask generated from RCNN segmentation D) DBSCAN based assignment of cells to a cluster.

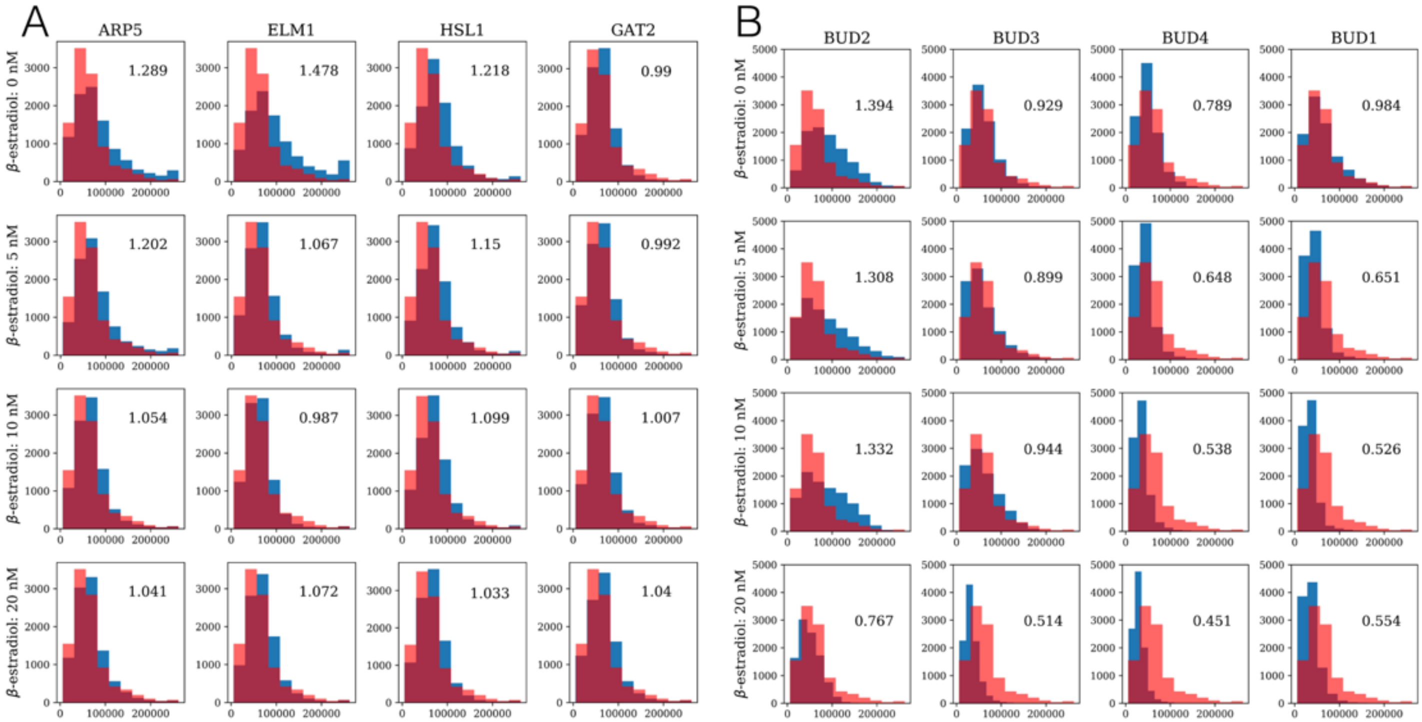


**Figure S5** A) Screen for increase in cluster size distribution using strains with β-estradiol inducible control of cell-shape properties. Size distribution of *ace2*Δ, with all other relevant genes in their wild-type form, shown in red and *ace2*Δ with inducible, β-estradiol control of the specific gene shown in blue. B) Screen for monotonic and significant increase in cluster size distribution using strains with inducible, β-estradiol control of genes that regulate the budding pattern. Size distribution of *ace2*Δ shown in red and *ace2*Δ with β-estradiol control of specific gene shown in blue. The number represents fold change in mean cluster size (measured as forward scatter area) with respect to size of *ace2*Δ cluster.

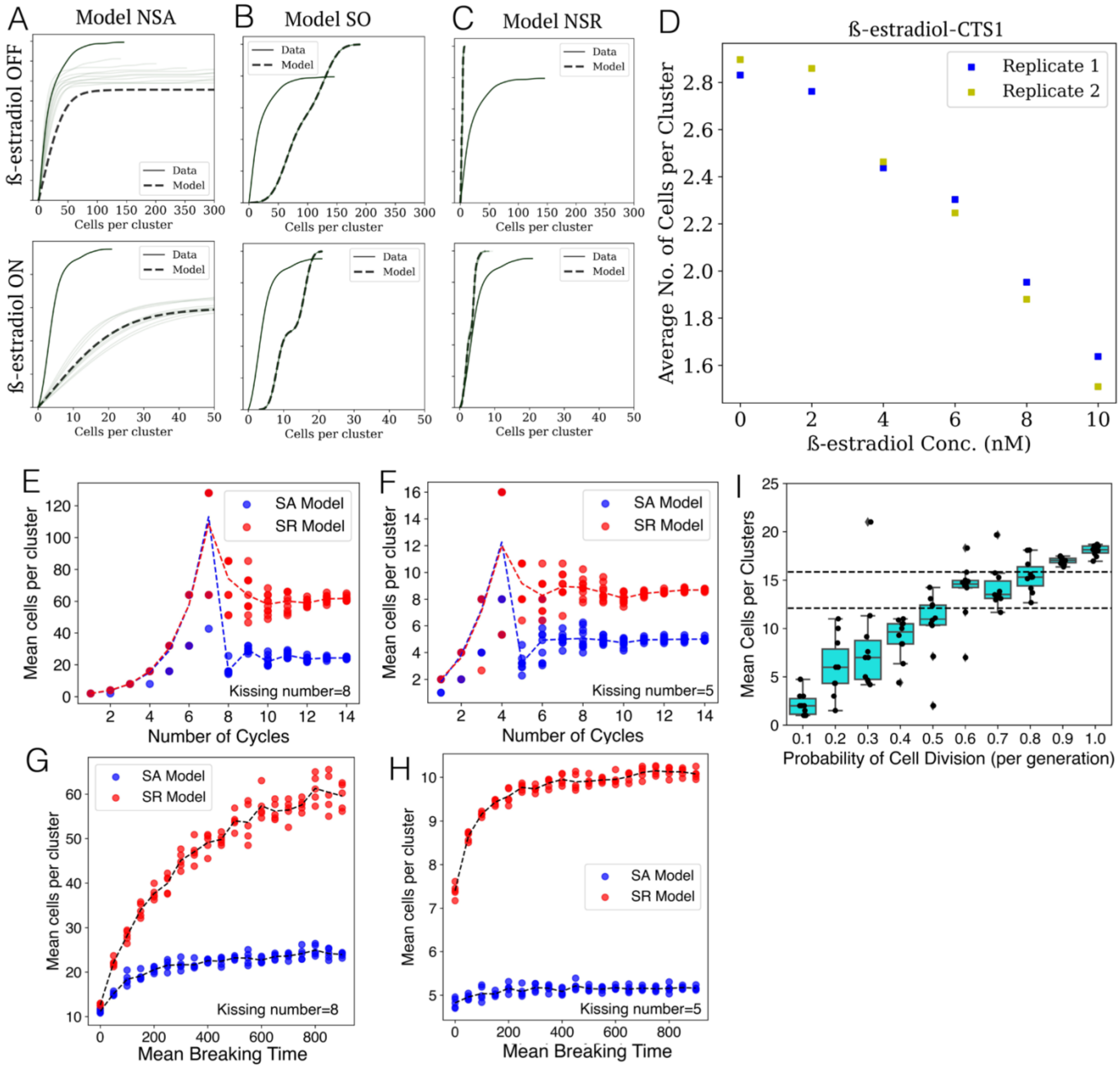


**Figure S6** Cluster size distribution generated from NSA (A), SO (B) and NSR (C) -based SA dynamic network models for high (β-estradiol OFF) and low (β-estradiol ON) kissing number using an *ace2*Δ strain with *BUD2* under the control of a β-estradiol inducible promoter. The gray lines represent independent simulation runs. D) Change in mean cluster size in an *ACE2* strain with *CTS1* under the control of a β-estradiol inducible promoter. E) and F). The effect of number of simulated growth cycles on mean number of cells per cluster for E) Kissing number ($\kappa$) = 8 and F) Kissing number ($\kappa$) = 5. G) and H). The effect of mean link-breaking time ($1/\delta$) on mean number of cells per cluster for G) Kissing number ($\kappa$) = 8 and H) Kissing number ($\kappa$) = 5. The dots represent the mean value of cluster size for independent simulations with the same parameters. I) Effect of asynchronous cell division on mean number of cells per cluster for $\kappa = 8$ using SA model. The probability of cell division determines how many cells are allowed to divide stochastically per generation. For example, a probability of 0.7 implies only 70% of the cells in the population divide and rest must wait for the next division event (after a unit division time). The dashed lines show mean number of cells per cluster from experiments for two replicates. The dots show independent simulations with different seed values.

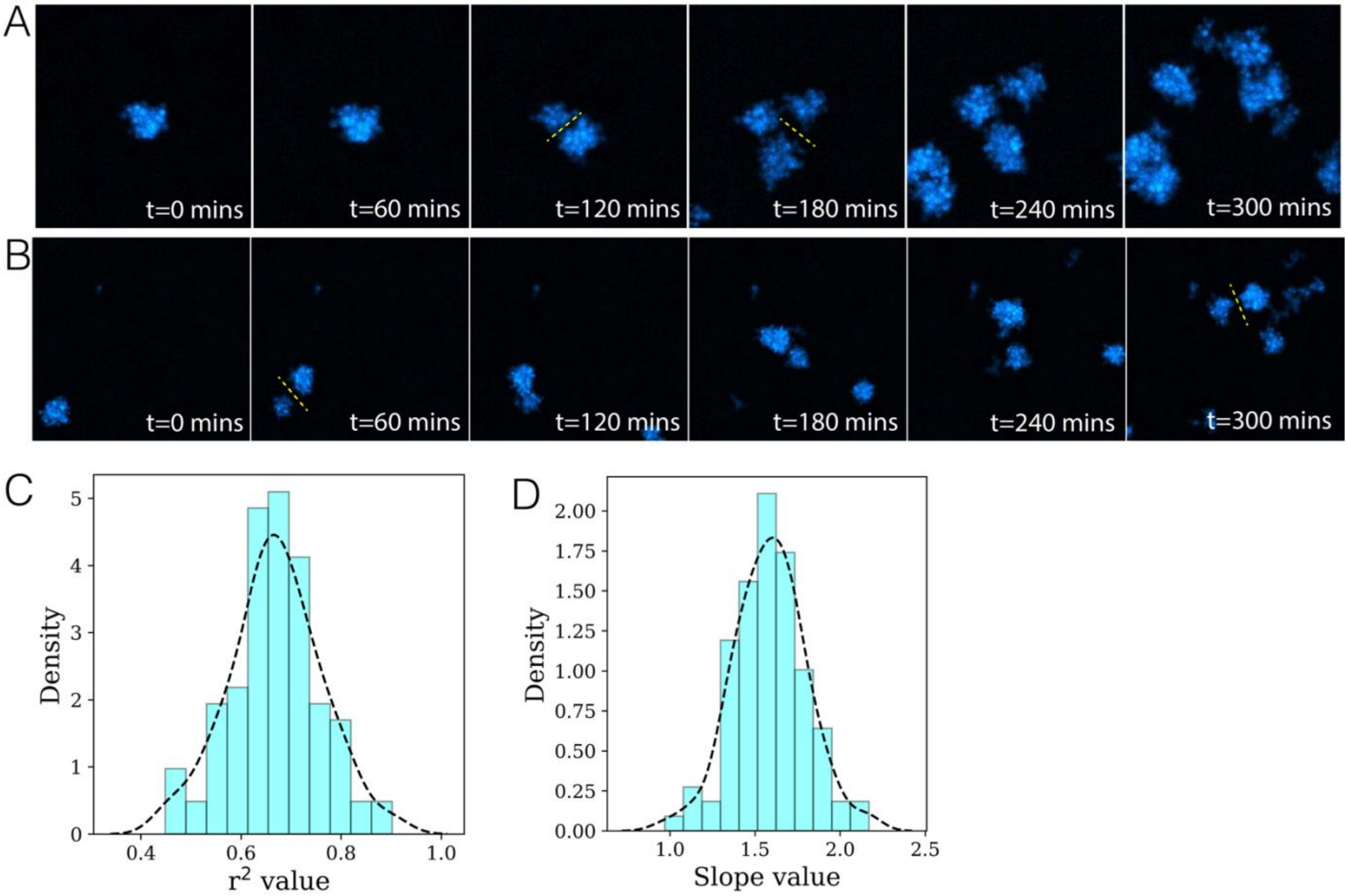


**Figure S7 (**A)-(B) Two independent time-traces of growth and fracture of multicellular clusters (C) $r^2$ value for comparison of model and experiment bin fraction for multiple rounds of independent simulations. (D) Slope values between experiment and simulation bin fraction. A value of 1 indicates perfect agreement. Bin fraction is defined as the fraction of total clusters that fall into a given cluster size range. The slope measures the correlation between mother: daughter cluster size ratios estimated from experiments and ratios obtained from simulations.

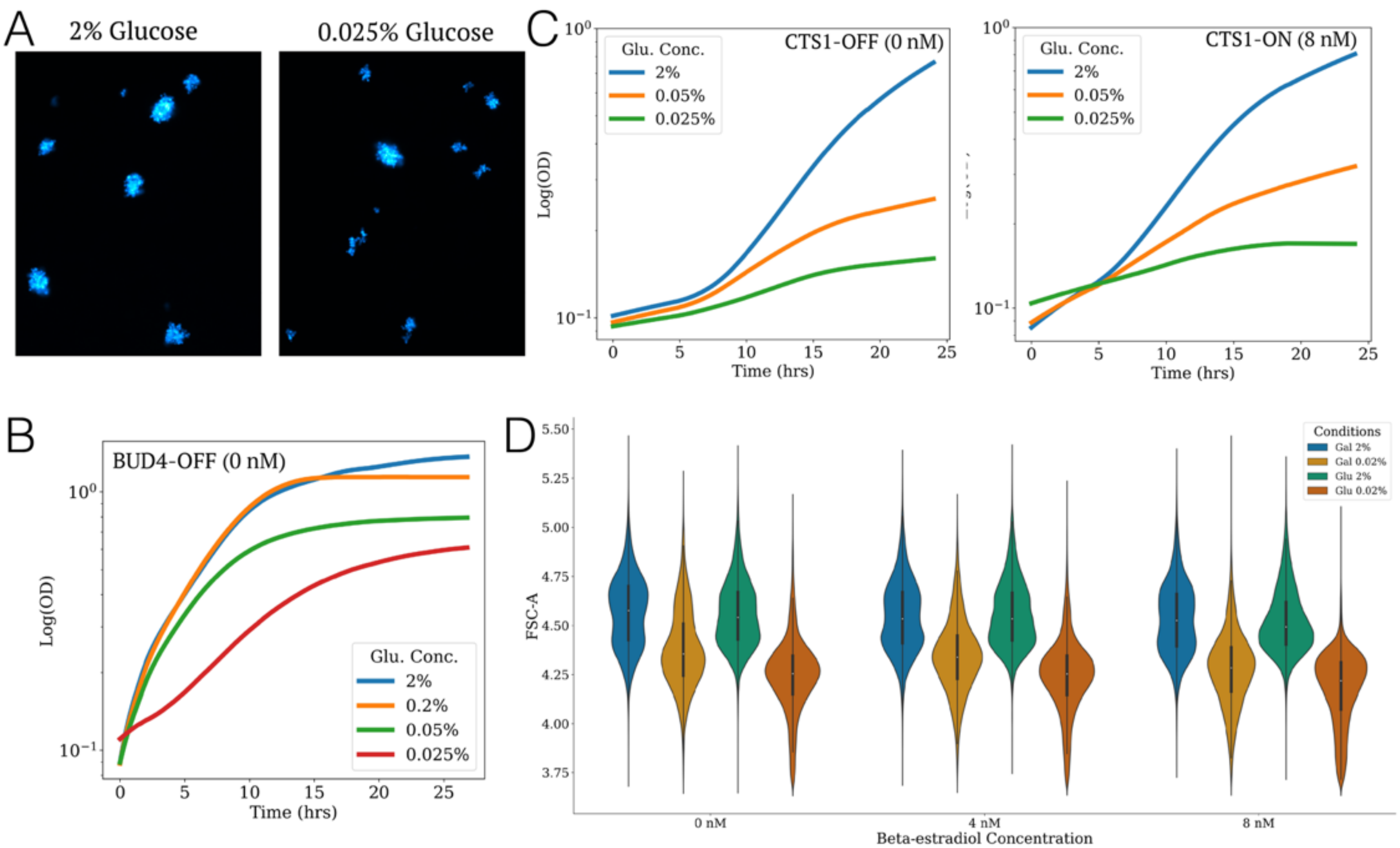


**Figure S8** A) Images of clusters with uninduced Bud4p in an *ace2*Δ strain, in which Bud4 is expressed from a β-estradiol-activated promoter, for two different glucose concentrations. The size distribution of these clusters is shown in Figure 4 B. B) Growth curves for cultures of the same strain. C) Growth curves for cultures of an *ACE2* strain with uninduced Cts1p expression and with partial (8 nM β-estradiol) induction of Cts1p D) Flow cytometry-based cluster size measurements for growth in two different concentrations of glucose and galactose and for various levels of Cts1p induction.

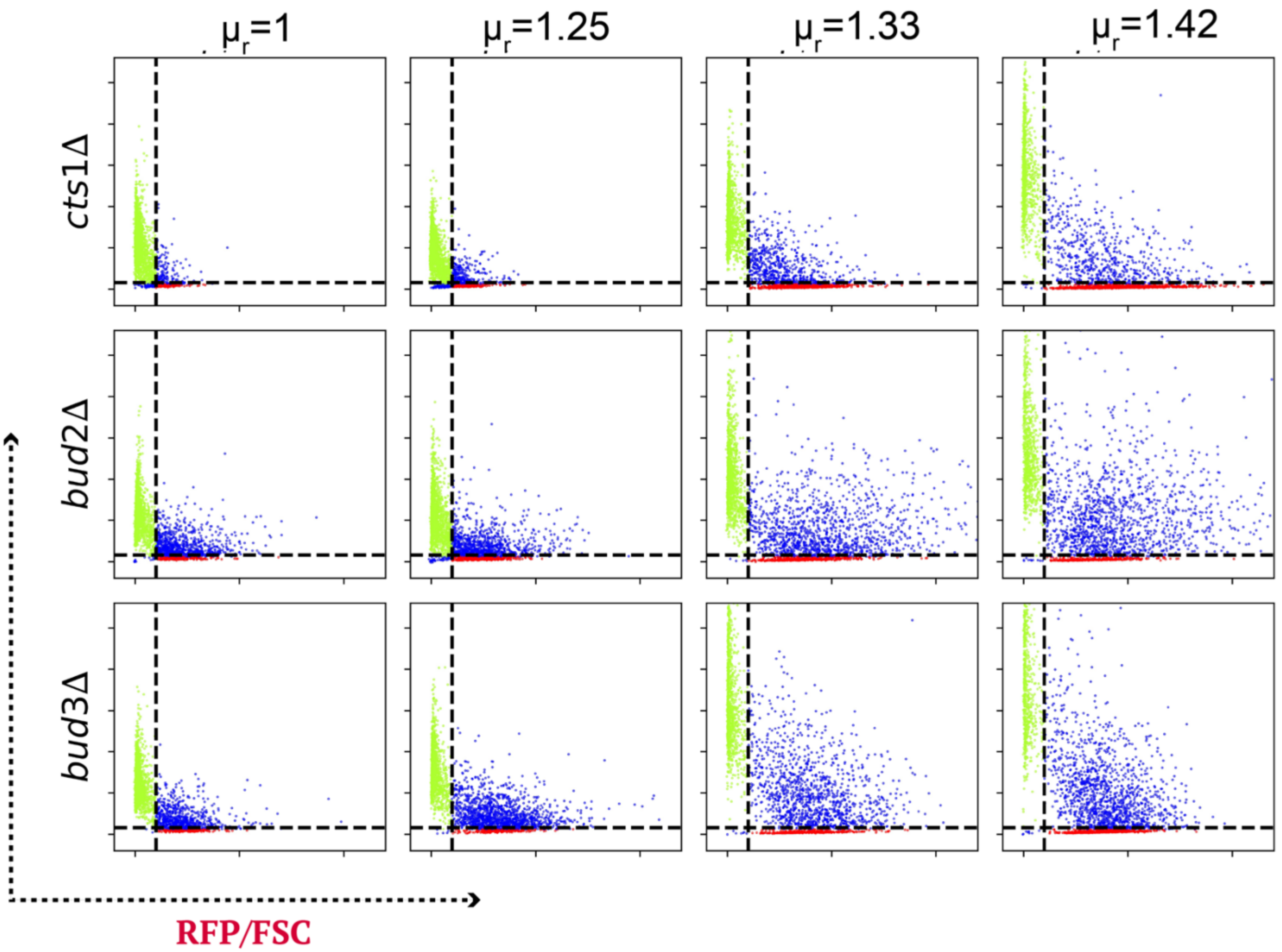


**Figure S9** Scatterplots for determining fraction of clusters comprised of purely somatic (green dots), purely germ (red dots), or mixed (germ and somatic, blue dots) cells in various backgrounds at four different concentrations of cycloheximide that produce an increasing growth advantage for germs cells. $\mu_r$ is the relative growth advantage of germ cells compared to somatic cells.

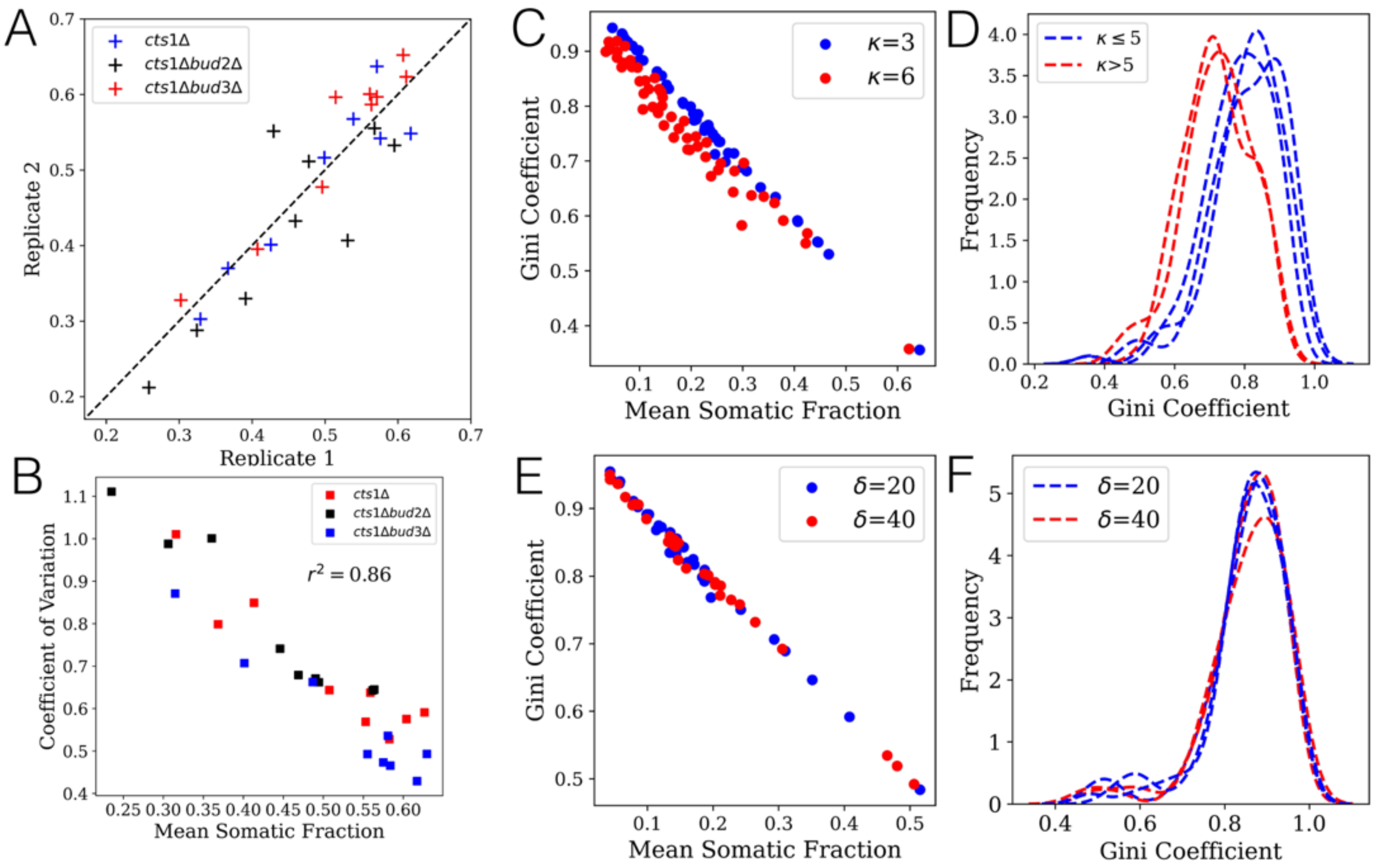


**Figure S10** A) Correlation between mean fraction of somatic cells in a cluster two independent biological replicates for three strains used for estimating cluster composition B) Relation between mean somatic fraction and its coefficient of variation for various strain backgrounds C) Predicted relation from SA model between mean somatic fraction and Gini coefficients for two kissing numbers. Dots show separate simulations where the net switching rate was varied D) Distribution of Gini coefficients for two different kissing number. E) Relation between mean somatic fraction and its coefficient of variation for various strain backgrounds for two different link breaking rates F) Distribution of Gini coefficients for two different link breaking rates.

**Table S1**

| Strain Name | Genotype |
|---|---|
| yPN073 | *can1-100 his3-11,15::PACT1(-1-520)-LexA-ER-haB42-TCYC1 BUD4-S288C prCTS1::PTEF-HygMX-TTEF-insul-(lexA-box)1-PminCYC1* |
| yPN078 | *can1-100 his3-11,15::PACT1(-1-520)-LexA-ER-haB42-TCYC1 BUD4-S288C prACE2::PTEF-HygMX-TTEF-insul-(lexA-box)1-PminCYC1* |
| yPN081 | *can1-100 his3-11,15::PACT1(-1-520)-LexA-ER-haB42-TCYC1 BUD4-S288C prHSL1::PTEF-HygMX-TTEF-insul-(lexA-box)1-PminCYC1* |
| yPN082 | *can1-100 his3-11,15::PACT1(-1-520)-LexA-ER-haB42-TCYC1 BUD4-S288C prARP5::PTEF-HygMX-TTEF-insul-(lexA-box)1-PminCYC1* |
| yPN083 | *can1-100 his3-11,15::PACT1(-1-520)-LexA-ER-haB42-TCYC1 BUD4-S288C prELM1::PTEF-HygMX-TTEF-insul-(lexA-box)1-PminCYC1* |
| yPN101 | *can1-100 his3-11,15::PACT1(-1-520)-LexA-ER-haB42-TCYC1 BUD4-S288C prBUD2::PTEF-HygMX-TTEF-insul-(lexA-box)1-PminCYC1* |
| yPN102 | *can1-100 his3-11,15::PACT1(-1-520)-LexA-ER-haB42-TCYC1 BUD4-S288C prBUD3::PTEF-HygMX-TTEF-insul-(lexA-box)1-PminCYC1* |
| yPN103 | *can1-100 his3-11,15::PACT1(-1-520)-LexA-ER-haB42-TCYC1 BUD4-S288C prBUD4::PTEF-HygMX-TTEF-insul-(lexA-box)1-PminCYC1* |
| yPN104 | *can1-100 his3-11,15::PACT1(-1-520)-LexA-ER-haB42-TCYC1 BUD4-S288C prBUD1::PTEF-HygMX-TTEF-insul-(lexA-box)1-PminCYC1* |
| yPN119 | *yMEW208* |
| yPN121 | *yMEW208 BUD2Δ::kanMX* |
| yPN125 | *yMEW208 BUD3Δ::kanMX* |
| yPN280 | *yPN103 HOΔ::prACT1-yCerulean-ADH1tr-KanMX* |
